# In Vivo Three-dimensional Brain Imaging with Chemiluminescence Probes in Alzheimer’s Disease Models

**DOI:** 10.1101/2023.07.02.547411

**Authors:** Jing Zhang, Carly Wickizer, Weihua Ding, Richard Van, Liuyue Yang, Biyue Zhu, Jun Yang, Can Zhang, Shiqian Shen, Yihan Shao, Chongzhao Ran

## Abstract

Optical three-dimensional (3D) molecular imaging is highly desirable for providing precise distribution of the target-of-interest in disease models. However, such 3D imaging is still far from wide applications in biomedical research; 3D brain optical molecular imaging, in particular, has rarely been reported. In this report, we designed chemiluminescence probes with high quantum yields (QY), relatively long emission wavelengths, and high signal-to-noise ratios (SNRs) to fulfill the requirements for 3D brain imaging in vivo. With assistance from density-function theory (DFT) computation, we designed ADLumin-Xs by locking up the rotation of the double-bond via fusing the furan ring to the phenyl ring. Our results showed that ADLumin-5 had a high quantum yield of chemiluminescence and could bind to amyloid beta (Aβ). Remarkably, ADLumin-5’s radiance intensity in brain areas could reach 4×10^7^ photon/s/cm^2^/sr, which is probably 100-fold higher than most chemiluminescence probes for in vivo imaging. Because of its strong emission, we demonstrated that ADLumin-5 could be used for in vivo 3D brain imaging in transgenic mouse models of Alzheimer’s disease (AD).

**Significance Statement:** Although MRI, PET, CT, and SPECT have been routinely used for 3D imaging, including 3D brain imaging, they are considerably expensive. Optical imaging is largely low-cost and high throughput. However, the 3D capacity of optical imaging is always limited. Obviously, optical 3D molecular imaging is highly challenging, particularly for 3D brain imaging. In this report, we provided the first example of 3D brain imaging with chemiluminescence probes ADLumin-Xs, which have advantages in quantum yields (QY), emission wavelengths, and signal-to-noise ratios (SNRs) to fulfill the requirements for 3D brain imaging. And we believe that such 3D capacity is potentially a game-changer for brain molecular imaging in preclinical studies.

## Introduction

Spatial information is essential in biomedical imaging to accurately diagnose diseases and understand fundamental biological mechanisms (1, 2, 3). Microscopy imaging has been widely used to provide spatial information via two-dimension (2D) and three-dimension (3D) imaging at the micro-scale of nanometers to micrometers (3, 4). At the macro- and meso-scale, MRI, PET, CT, and SPECT have been routinely used to provide spatial information *via* 3D reconstruction imaging. Comparatively, 3D optical imaging with molecular probes, including fluorescence imaging, bio-luminescence imaging, chemiluminescence imaging, and Cerenkov luminescence imaging, has yet to be widely applied (5).

Although 3D optical fluorescence imaging and bioluminescence imaging have been explored for tumor imaging (6); 3D optical brain imaging has been rarely reported (7, 8, 9). For tumor imaging, due to the concentrated mass of a tumor, the signal quickly reaches the threshold required for 3D reconstruction. For 3D non-tumor brain imaging, it has been believed to be more challenging, because several hurdles need to be overcome: 1) the strong scattering by brain skulls (10), 2) the mass/distribution of the targets are normally dispersed, and 3) the strong light attenuation by the brain tissue and skull (11).

Although numerous attempts, including efforts from our group, have been made to achieve reliable 3D optical brain imaging, there has been limited success so far (9). Multiple factors have profound influences on 3D optical brain imaging, including: 1) signal strength for the 3D reconstruction, 2) light penetration capacity to reach the targets in a brain, and 3) signal-to-noise ratio (SNR) or signal-to-background ratio (SBR) for providing high-quality signals for the 3D reconstruction. For fluorescence 3D imaging, low SNRs have severely hampered the 3D reconstruction (12), and the low SNRs are partially due to the two-way (in-and-out) traveling of photons that are involved in excitation and emission (13). Given that chemiluminescence is one-way traveling of the emitted photons, chemiluminescence imaging could provide considerably high SNRs (14). Based on these facts, we speculated that 3D chemiluminescence brain imaging could be feasible if the probes meet the following requirements: 1) strong signals from the brain, 2) excellent penetration of the emitted light, and c) high SNR in the interested brain areas.

In our previous studies, we reported that ADLumin-1 is a turn-on chemiluminescence probe for amyloid beta (Aβ) species in Alzheimer’s disease (AD) (15). In comparison to other chemiluminescence probes that rely on ROS or enzymatic triggering for photon emission, ADLumin-1 offers the advantage of utilizing oxygen to initiate photon emission. In addition, the chemiluminescence of ADLumin-1 can be further dramatically enhanced in the presence of Aβ aggregates. However, a short emission wavelength and a low quantum yield are the two shortcomings associated with ADLumin-1. In this report, we designed a series of chemiluminescence probes ADLumin-Xs (X = 5-8) to achieve high quantum yields and/or longer emission wavelengths to meet the requirements for 3D optical brain imaging (Fig. 1). Indeed, we have successfully performed in vivo 3D brain imaging with ADLumin-5 in mice and differentiated wild type (WT) and AD mice. To the best of our knowledge, our in vivo 3D brain imaging is the first of its kind with chemiluminescence probes for brain tomography imaging.

**Figure 1.**
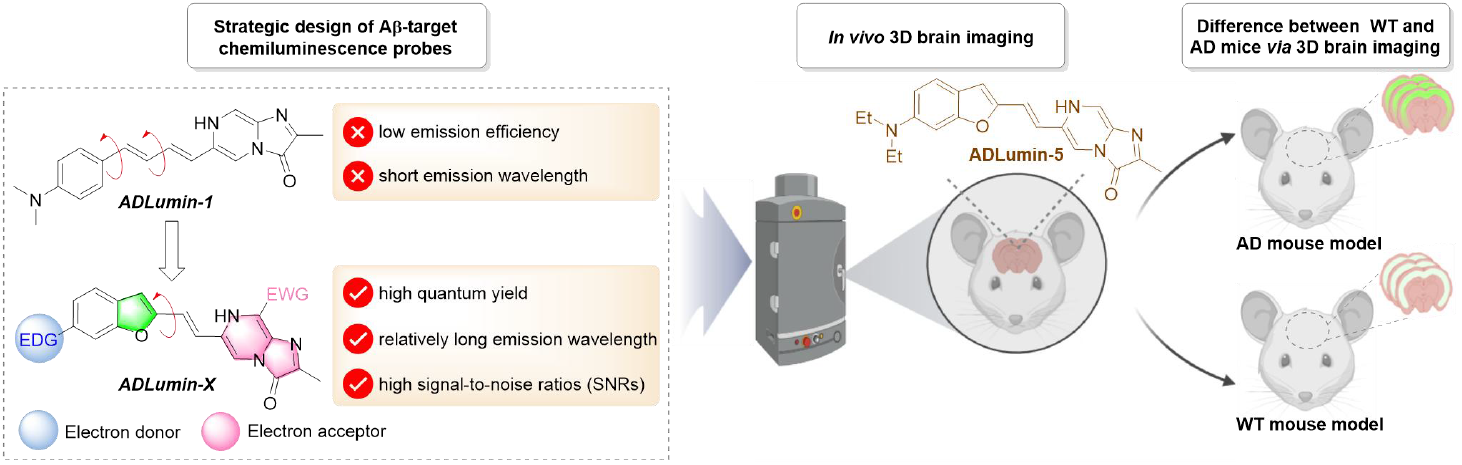
The design strategy of ADLumin-Xs probes and illustration of this work for in vivo 3D imaging. In this work, we first validated that the constraining of the double-bond rotation could result in significant improvement of quantum yields, and the probes were comprehensively tested in vitro for their features, including spectral studies, exploratory mechanism investigation, and Aβ binding. Lastly, we demonstrated that 3D brain tomography imaging is feasible for in vivo studies.

## Results

### 1. Design of ADLumin-Xs (X = 5-8) and density-functional theory (DFT) computation

In ADLumin-1, the chemiluminescent moiety imidazo[1,2-a]<u>p</u>yrazin-3(7H)-<u>o</u>ne (IPO) is π-conjugated *via* double bonds with the electron-donating phenyl ring. We speculated that the rotation of the conjugated double bonds in ADLumin-1 caused its low QY, due to non-radiative decay (Fig. 1). To overcome the weakness of low QY of ADLumin-1, we proposed to constrain the rotation of the double-bond connecting to the phenyl ring *via* fusing with the five-membered furan ring (X = 5). Meanwhile, the fusion of the furan ring is also expected to increase the electron density of the whole system, which can narrow the HOMO-LUMO gap to enable the emission at longer wavelengths. To further extend the emission of ADLumin-5, we reasoned that introducing an electron-withdrawing cyano-group at the IPO moiety could facilitate electron delocalization from the donor moiety (benzofuran) to the acceptor (IPO). In this regard, ADLumin-6 was designed. To investigate the influence of the *N,N’*-dialkylamino-group on the phenyl ring, ADLumin-7 and -8 were designed by replacing the *N,N’*-dimethylamino-group with -OH and -OMe, respectively (Fig. 2a).

**Figure 2.**
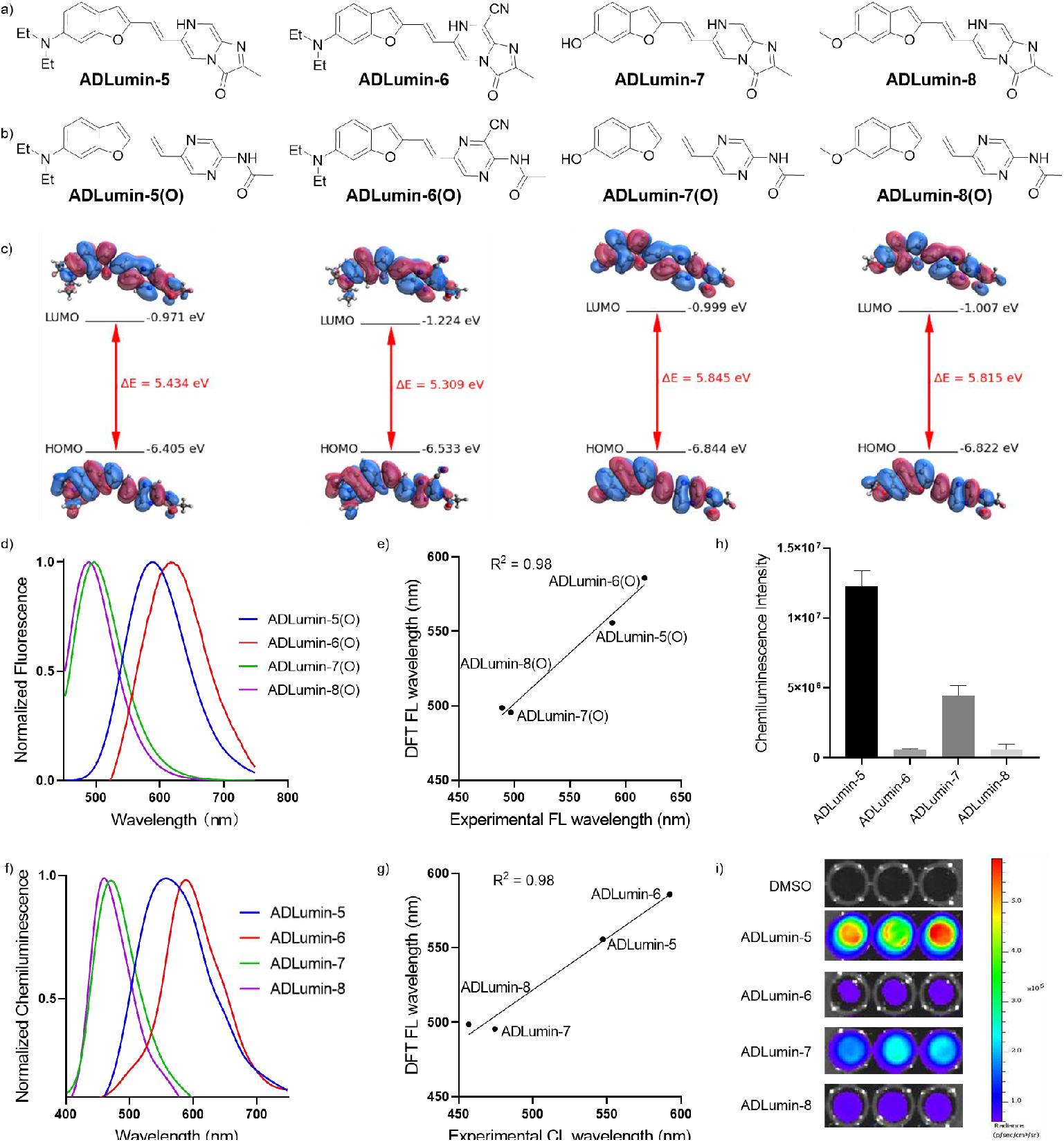
DFT computation and characterization of ADLumin-Xs. a) The structure of ADLumin-Xs; b) The structure of ADLumin-Xs(O); c) HOMO, LUMO, and HOMO-LUMO energy gaps for ADLumin-Xs(O) in DMSO calculated at optimized excited state geometries with wB97X-D functional and 6-31+G* basis; d) Fluorescence emission spectra of ADLumin-Xs(O) in DMSO; e) Correlation of fluorescence emission wavelength between experimental data and DFT data (R^2^ = 0.98); f) Chemiluminescence spectra of ADLumin-Xs in DMSO; g) Correlation of chemiluminescence emission wavelength between experimental data and DFT data (R^2^ = 0.98); h) Chemiluminescence intensity of ADLumin-Xs in DMSO; i) In vitro chemiluminescence images of ADLumin-Xs in DMSO.

To verify our design rationale, TD-DFT calculations were carried out at the ωB97X-D/6-31+G* level (16). Because chemiluminescence emission was generated from the high energy (excited) intermediate of the open-ring products ADLumin-Xs(O)* (Fig. 2b), we compared the DFT calculated emission wavelengths of ADLumin-Xs(O) with the measured fluorescence/chemiluminescence emission wavelengths and proved the rationality by whether the trend was consistent. As depicted in Figure 2c, the HOMO and LUMO were calculated at excited state geometries for ADLumin-Xs(O). The energy gap between LUMO and HOMO of ADLumin-5(O) was about 5.434 eV, which was lower than ADLumin-7(O) and -8(O) (5.845 eV and 5.815 eV, respectively). As we expected, the electron-withdrawing cyano-group led ADLumin-6(O) to show a smaller HOMO-LUMO gap, which suggests a slightly longer emission wavelength (Fig. 2c).

We measured their fluorescence emission spectra in DMSO (Fig. 2d), and found that the trend of theoretical prediction was consistent with the experimental fluorescence emission for all probes (Fig. 2e). Specifically, the introduction of -CN group extended the emission wavelengths, while replacing the diethylamino group with -OH and -OMe shortened the emission wavelengths. We also recorded their chemiluminescence spectra in DMSO, and found that the emission peaks were consistent with the computed fluorescence emission wavelengths that are based on ADLumin-Xs(O), the open-ring products of the photon emitting reaction (Fig. 2f, 2g). Among the probes, ADLumin-6 showed the longest emission wavelength. Similar to the fluorescence spectral studies, ADLumin-7 and -8 showed shorter chemiluminescence emission wavelengths, again suggesting that the diethylamino group in ADLumin-5 and -6 are needed for longer emission wavelengths. Taken together, our computation results confirmed the rationale of our probe design.

### 2. Characterization of ADLumin-Xs (X= -5, -6, -7, and -8)

We found that all the ADLumin-Xs at 2.5 μM could generate strong chemiluminescence in DMSO solutions (Fig. 2h, 2i) and high signal-to-noise ratio (SNRs > 600), but no significant signal in other organic solvents such as acetone, methanol, and methylene chloride (SI Fig. S1). We also observed intensity enhancement if pure O_2_ gas was bubbled into the DMSO solutions, suggesting that ADLumin-Xs were O_2_-sensitive (SI Fig. S2). In addition, we observed that ADLumin-Xs could produce weak chemiluminescence in PBS buffer solutions (SI Fig. S3), and SNRs were about 100. However, the intensities were 130-fold lower than the intensity from the DMSO solutions (ADLumin-5 as the example). This is probably due to the formation of dipole-dipole moments between water molecules and the high-energy intermediates that can significantly reduce the radiative decay of the HEIs, and, consequently, low photon emitting efficiency in the PBS buffer.

### 3. Mechanism of photon emitting

Although the auto-oxidation of cypridina luciferin by O_2_ is well-known, the mechanism is still unclear. We used ADLumin-5 as an example to investigate the mechanism of the reaction. The IPO moiety in ADLumin-Xs is the core structural component of cypridina luciferin (17). Reportedly, the rate of autooxidation of cypridina luciferin analogues, such as CLA, varies with pH of the solutions (18, 19). To investigate the property changes of ADLumin-5 under different pH conditions, we conducted pH titration in buffers. We first used the absorption changes of ADLumin-5 under different pH conditions to calculate pKa values, and found that the pKa of ADLumin-5 was 7.56 (SI Fig. S4). Next, we monitored the changes in chemiluminescence intensities with the increasing pH. Similar to other cypridina luciferin analogues, the chemiluminescence intensity of ADLumin-5 showed apparent pH dependency (Fig. 3a and SI Fig. S5 for other probes). Notably, we observed strong chemiluminescence under low pH conditions (pH = 2-5) (Fig. 3a, 3b). In the 1960s and 1993, Goto and Fujimori et al. speculated that chemiluminescence would be observable under low pH conditions; however, they failed to observe the emission of chemiluminescence because the intensity of the chemiluminescence emission was so weak in the autoxidation of CLA (19). Clearly, our results provided strong evidence for validating their hypothesis. And on the other hand, it also confirmed that the chemiluminescence intensity of ADLumin-5 was strong enough to be detected at low pH conditions. Furthermore, we evaluated the emission wavelength of ADLumin-5 under different pH conditions and observed that it had a shorter emission wavelength at low pH conditions and a longer wavelength at high pH conditions (SI Fig. S6). It suggested that it had two pathway to emit the light: 1) the excited states of neutral forms, which was protonated at low pH (ADLumin-5(O)* as an example) can emit short wavelength light; 2) the excited states of anionic forms can emit long wavelength light at relatively high pH (ADLumin-5(O)*^−^ as an example) (Fig. 3e).

**Figure 3.**
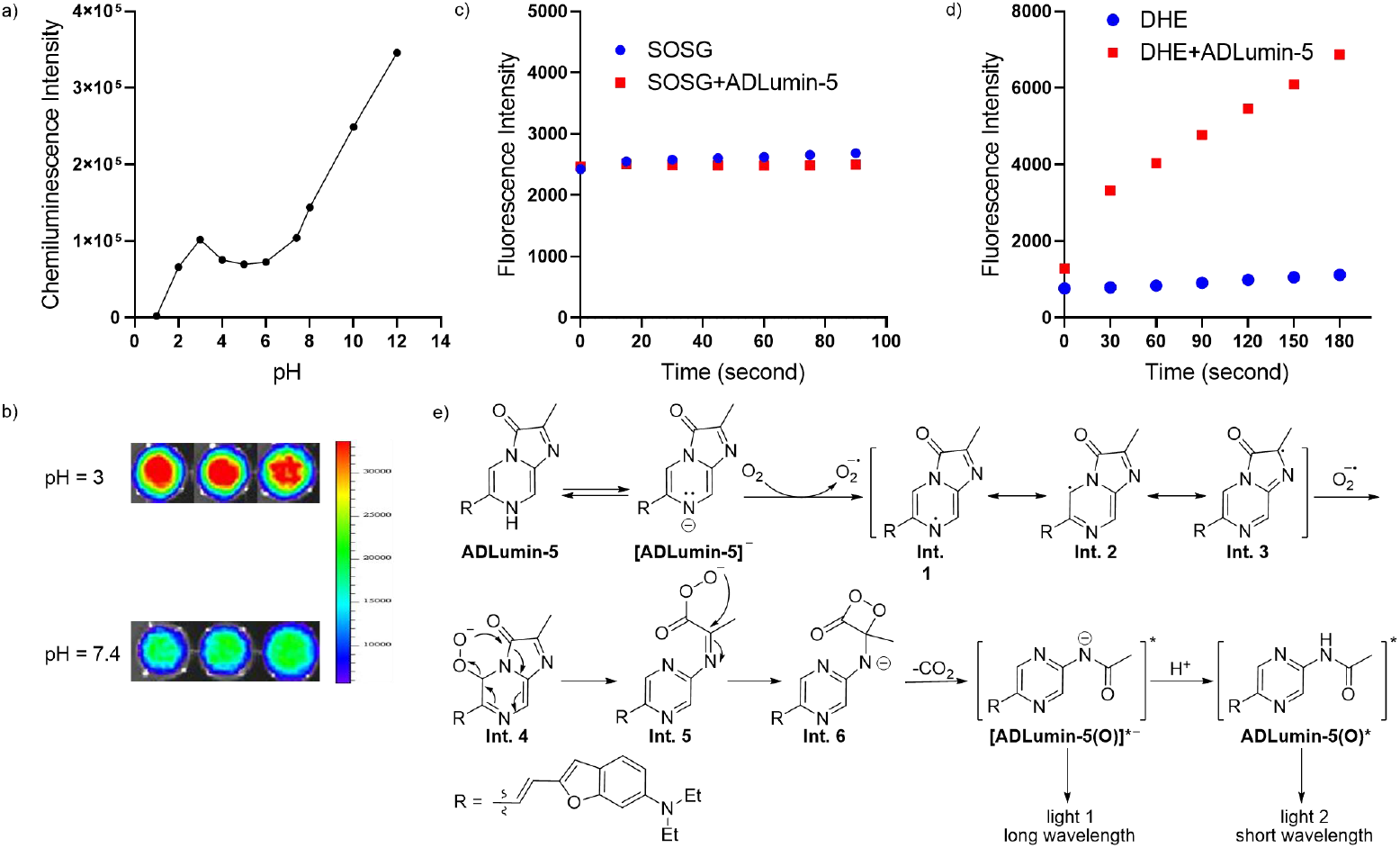
Mechanism study of ADLumin-Xs. a) The pH dependence of ADLumin-5 in solutions; b) Representative images at pH 3 and 7.4 for ADLumin-5; c) The detection of singlet oxygen (^1^O_2_) with SOSG during the chemiluminescence generation from ADLumin-5. No significant amount of ^1^O_2_ was detected in the solutions of ADLumin-5; d) The detection of superoxide anion radical (O_2_^−•^) detection by DHE during the chemiluminescence generation from ADLumin-5. Significant amounts of O_2_^−•^ were detected in the solutions of ADLumin-5; e) The proposed mechanism of chemiluminescence emission from ADLumin-Xs.

Based on the evidence that superoxide dismutases (SOD) can scavenge superoxide anion radical (O_2_^−•^), Goto et al. speculated that the key oxygen species involved in the reaction of cypridina luciferin was O_2_^−•^, instead of singlet oxygen (^1^O_2_) (20). Since ADLumin-Xs also contain the key IPO moiety of cypridina luciferin, it is interesting to investigate whether ADLumin-5 also follows a similar reaction mechanism. To this end, we first measured the chemiluminescence intensity changes before and after the addition of SOD; however, only a slight decrease in intensity was observed (SI Fig. S7). To clarify the mechanism further, we measured the levels of O_2_^−•^ using dihydroethidium (DHE) to record its fluorescence changes (Fig. 3c) (21). Indeed, we observed a significant DHE fluorescence intensity increase with ADLumin-5. In contrast, no increase was observed if no ADLumin-5 was added, confirming that O_2_ ^−•^ is the key oxygen species to initiate the auto-oxidation. It is likely that no significant amount of ^1^O_2_ was generated during the auto-oxidation of ADLumin-5. We further excluded the involvement of ^1^O_2_ via measuring the ^1^O_2_ level with a singlet oxygen sensor green (SOSG), evidenced by no significant difference observed with and without ADLumin-5 (Fig. 3d).

Based on the above data, we proposed a light-emitting mechanism of ADLumin-5. Firstly, ADLumin-5 was deprotonated to form [ADLumin-5]^−^ under a buffer solution. Through single-electron transfer (SET), oxygen (triplet) was activated and converted into superoxide anion radical (O ^−•^) by accepting the electron from [ADLumin-5]^−^. The active O ^−•^ then attacked the IPO moiety to form intermediate 4 (Int. 4) and occurred a series of intramolecular electron transfer to form Int.6, which was easy to release carbon dioxide. Finally, the two forms of high-energy intermediates returned back to the ground state through photon emitting (Fig. 3e). Although more detailed studies are needed for the mechanism investigation, our results are very important for the validation of the mechanisms that have been proposed several decades ago (19).

### 4. ADLumin-X’s binding to Aβ aggregates

We evaluated the binding of ADLumin-Xs with Aβ_40_ aggregates in PBS buffer solutions. As expected, the chemiluminescence intensities of all the probes increased upon mixing with the Aβ aggregates (Fig. 4a). ADLumin-5 provided 18-fold amplification of the signal, and ADLumin-6 afforded the largest amplification folds (118-fold). Interestingly, although ADLumin-6 showed the largest amplification, its absolute intensity is relatively low (Fig. 4b). Among the probes, ADLumin-5 provided the highest absolute intensity. The signal amplification of ADLumin-Xs can be ascribed to their binding to the hydrophobic environment inside Aβ fibrils and the enhanced conformational rigidity of the probes upon Aβ binding (22).

**Figure 4.**
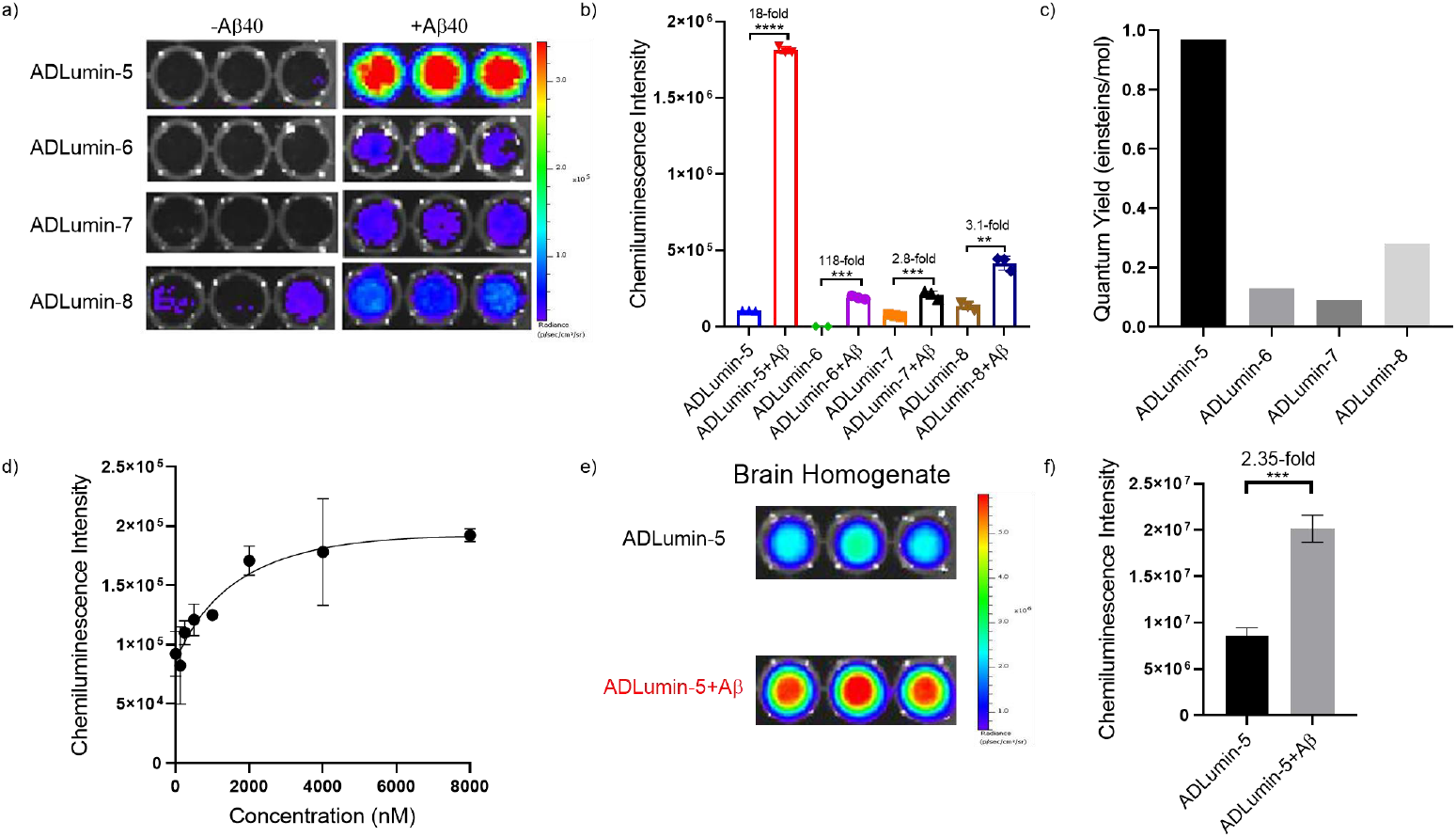
In vitro characterization of chemiluminescence properties of ADLumin-Xs in the presence of Aβs. a) In vitro chemiluminescence images of ADLumin-Xs with/without Aβ_40_ aggregates in PBS; b) Quantification of chemiluminescence intensity in a); c) Quantum yield of ADLumin-Xs with Aβ_40_ aggregates in PBS; d) Binding affinity assay between Aβ_40_ aggregates and ADLumin-5. Data point with its error bar indicates mean ± s.d. derived from n=3 independent samples; e) In vitro chemiluminescence images of ADLumin-5 with/without Aβ_40_ aggregates in mouse brain homogenate; f) Quantification of chemiluminescence intensity in e).

To further confirm the excellent brightness of ADLumin-5 in the presence of Aβ aggregates, we measured the quantum yields of the probes by using luminol as the reference (23). As expected, ADLumin-5 showed the highest QY (0.97), compared to other probes (QY= 0.13, 0.09, 0.28 for ADLumin-6, -7, and -8, respectively) (Fig. 4c).

We also measured the binding constant Kd of ADLumin-Xs with different concentrations of Aβ_40_ aggregates *via* titration and found that Kd was 2.1 μM for ADLumin-5, and Kd = 28.8, 0.21, and 0.53 μM for ADLumin-6, -7, and -8, respectively (Fig. 4d and SI Fig. S8). From the titration curve, it is clear that the intensity increase of ADLumin-5 is concentration-dependent in the range of 0-2.0μM. To investigate whether ADLumin-5 can be used under biologically relevant environments, we incubated ADLumin-5 with Aβ_40_ aggregates in the presence of mouse brain homogenates and found a 2.35-fold intensity increase in the Aβ_40_ group (Fig. 4e, 4f). Our results suggested that ADLumin-5 could be used for further in vivo imaging studies.

Given that ADLumin-5 and -6 have relatively longer emission wavelengths. In comparison, ADLumin-7 and -8 have shorter emission wavelengths, in the following studies, ADLumin-5 and -6 were selected for more intensive investigations on in vitro and in vivo imaging.

### 5. Mimic tissue penetration studies

Since tissue penetration is a key issue for in vivo optical imaging, we first evaluated ADLumin-Xs’ penetrating capacity under mimic conditions with chicken breast tissues. ADLumin-5 and -6 showed excellent penetration, about 2% light can be detected after penetrating 1.0 cm of the tissue (SI Fig. S9). To further confirm the penetration capacity, we used nude mice as the mimics by placing the probes under the belly and collected signals from the dorsal side (Fig. 5a). Similar to the results from the chicken tissue, we found that about 2% of light penetrated through the whole mouse body with ADLumin-5, and -6 (Fig. 5b).

**Figure 5.**
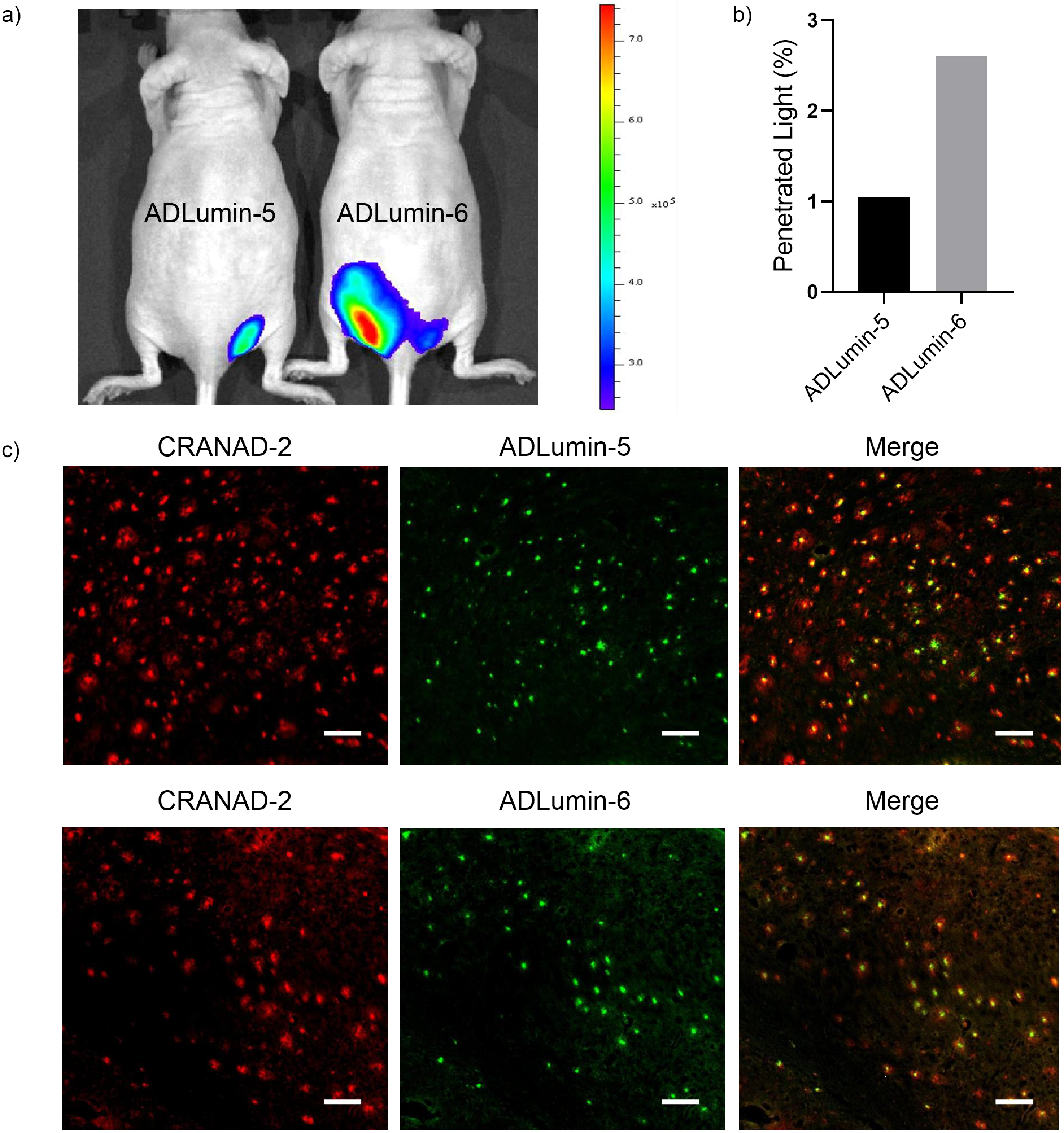
Chemiluminescence light penetration and tissue staining. a) Representative images that were captured from the dorsal side with an IVIS imaging system after placing the tubes filled with a solution of probes under the belly of the mice; b) Quantification of chemiluminescence intensity in a); c) In vitro brain slice imaging with ADLumin-5 and -6. Scale bar: 100 μm. The red spots indicate that plaques are stained by CRANAD-2, and the green spots indicate that plaques are stained by ADLumin-5 and ADLumin-6, respectively. Excellent co-localization could be observed from the staining with CRANAD-2 and ADLumin-5, and ADLumin-6.

### 6. AD mouse brain slide staining

Given that the binding target of ADLumin-Xs is aggregated Aβs, such as Aβ plaques, it is necessary to confirm whether ADLumin-Xs can bind to Aβ plaques in brain slides. In this regard, we incubated ADLumin-5 and-6 with brain slices from an 18-month-old 5xFAD mouse, and CRANAD-2, a NIRF probe from our group (24), was used as the positive control for plaque staining. Thioflavin T was not used because its emission wavelength is similar to ADLumin-Xs. As expected, ADLumin-5 and-6 showed an excellent staining capacity for Aβ plaques and excellent co-localization with CRANAD-2 in the mouse brain slides, further supporting that they are good candidates for in vivo imaging investigations (Fig. 5c).

### 7. In vivo 2D imaging with AD mouse models

Due to their relatively longer emission wavelengths, ADLumin-5 and -6 were selected for this in vivo imaging study. To investigate whether ADLumin-5 can be used for in vivo imaging to report the difference between wild-type (WT) and transgenic AD mice, we performed in vivo imaging on an IVIS imaging system with 6-month-old 5xFAD mice and age-matched WT mice. As expected, ADLumin-5 showed significantly higher chemiluminescence signals from the brain areas in 5xFAD mice, and the difference was 1.5-fold higher at 30 minutes post-injection of ADLumin-5 (Fig. 6a and SI Fig. S10a). Meanwhile, with the same imaging protocol, we found that ADLumin-6 could provide 5.1-fold higher signals from the AD group (Fig. 6b and SI Fig. S10b). The larger differences between the AD and WT groups are likely due to the higher signal amplification of ADLumin-6 with Aβs, which is consistent with the results from the in vitro solution test.

**Figure 6.**
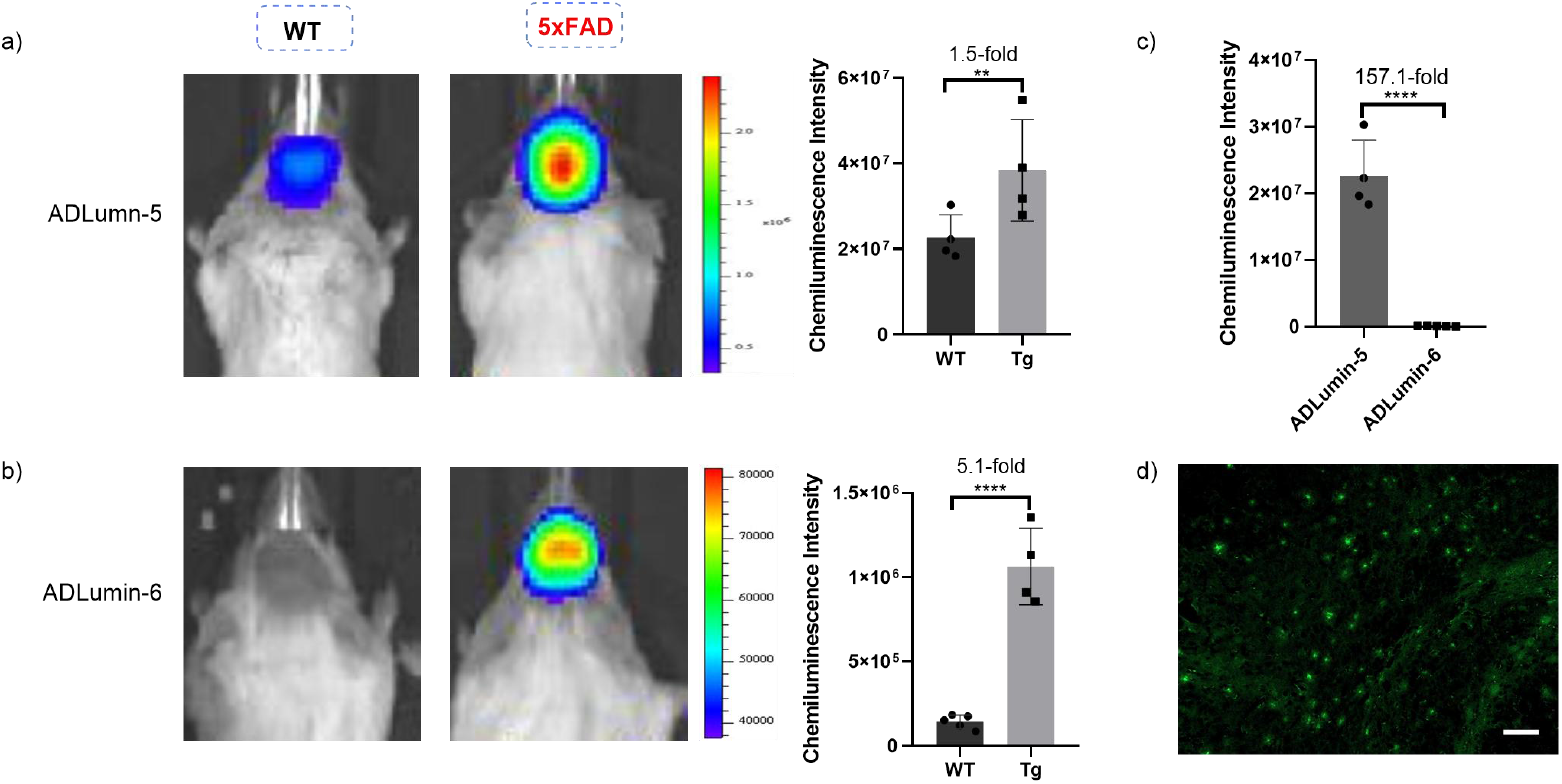
In vivo chemiluminescence imaging with ADLumin-5 and -6. a) Representative images of in vivo brain imaging and quantification of WT and 5xFAD mice with i.p. injection of ADLumin-5; b) Representative images of in vivo brain imaging and quantification of WT and 5xFAD mice with i.p. injection of ADLumin-6; c) Comparison of chemiluminescence intensities between ADLumin-5 and ADLumin-6 with WT mice, and 157-fold higher signals could be observed from ADLumin-5, suggesting ADLumin-5 could meet the requirements for 3D brain imaging; d) Ex vivo brain slice imaging with ADLumin-5. Scale bar: 100 μm. Error bar: Stdev; Data points with its error bar stands for mean ± s.d. derived from n = 4 independent animals. ****P-value < 0.0001; **P-value < 0.01.

Notably, ADLumin-5 and -6 showed very large differences in the chemiluminescence intensity from the brain areas of WT mice (Fig. 6c). ADLumin-5 had 157-fold higher signals, compared to ADLumin-6. Again, this data suggested that ADLumin-5 was highly bright, and the signal intensity (radiance of 4×10^7^ photon/s/cm^2^/sr) from this in vivo imaging was beyond the most reported chemiluminescence probes (mostly, the radiance range is in ∼10^5^ photon/s/cm^2^/sr).

To further confirm ADLumin-5’s in vivo labeling capacity, we performed ex vivo microscopic imaging with brain slides from the 5xFAD mice injected with ADLumin-5. Indeed, the plaques could be clearly visualized (Fig. 6d).

### 8. 3D imaging with ADLumin-5

The majority of 3D tomography optical imaging has been performed for tumor studies, likely this is due to the fact that tumors always have concentrated mass that can provide strong signals for 3D reconstruction. For tumors or other mass-concentrated targets, such as brown adipose tissue, 3D fluorescence, bioluminescence, chemiluminescence, and Cerenkov luminescence imaging have been reported (6, 25); however, rare examples could be found for non-tumor 3D brain imaging, such as 3D imaging of neurodegenerative diseases. Compared to other organs, brain imaging has an extra challenge, including skull scattering and low concentrations of target-of-interest.

In our studies, we used the diffuse luminescence tomographic algorithm (DLIT™) to estimate the 3D location and the photon flux of the sources in mouse brains (26). In this algorithm, the penetration capabilities of different wavelengths are used to compute the source locations and intensities *via* multiple spectral imaging. Specifically, in our studies, we collected chemiluminescence signals with 540-, 560-, 580-, 600-, 620-, 640-, 660-, 680- and 700-nm filters, and a signal threshold < 10.0% was used for the reconstruction. With ADLumin-5, we injected a 5xFAD mouse via tail vein at a dose of 4.0 mg/kg and started to collect the multiple spectral images 10 minutes post the injection. The acquisition time was 60 seconds for each filter, and the radiance from each filter was > 1*10^5^ photon/s/cm^2^/sr (SI Fig. S11). Upon completing the image collection, we performed 3D reconstruction with the DLIT algorithm. The reconstructed images are generally consistent with the distribution of Aβ deposits in brains, i.e., mainly in the cortex areas (Fig. 7a-c). Notably, our results suggested that the reconstructed depth was beyond 0.5 cm, which is quite impressive. Remarkably, the light source from the deep brain area also can be reconstructed (Fig. 7b, 7c, 7g). To the best of our knowledge, this 3D tomography is likely the first of its kind of optical brain imaging.

**Figure 7.**
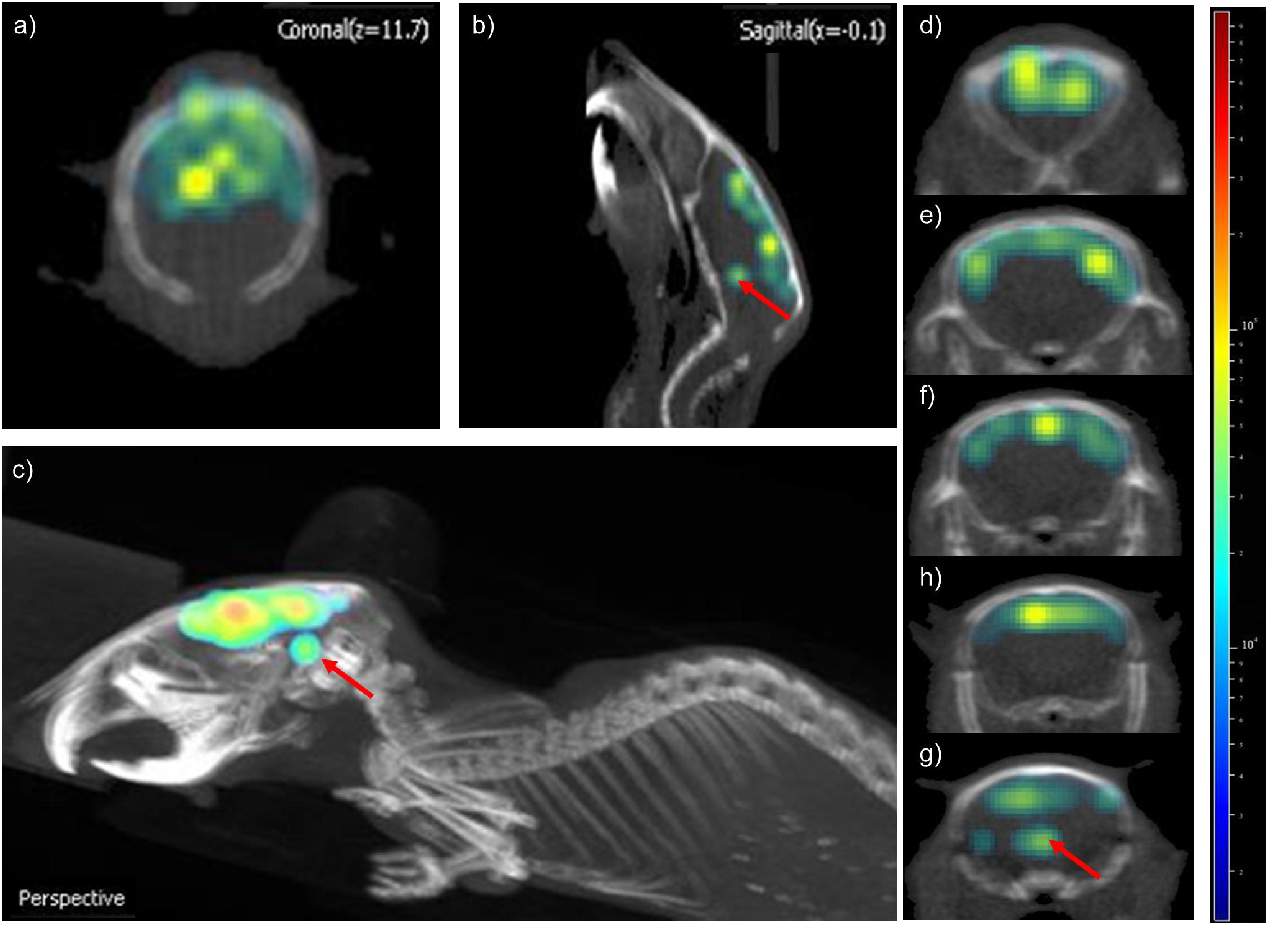
3D brain tomography imaging with ADLumin-5. a) Representative image of coronal view; b) Representative image of sagittal view; c) Representative reconstructed 3D brain image; d-g) Representative images of transaxial view. The red arrow shows the signal of the mouse brain at ∼ 0.5 cm depth, suggesting ADLumin-5 has an excellent penetration depth in vivo.

However, we failed to perform 3D imaging with ADLumin-6 by following the same imaging protocol. This is very likely due to the low intensity of ADLumin-6 from the brain areas, and this result also suggested that high intensities of a probe are critical for successful 3D brain imaging.

To investigate whether 3D brain tomography imaging can be used to report the difference between WT and AD mice, we performed 3D imaging with paired WT and AD mice simultaneously and with the same imaging protocol. From Fig. 8a, the difference between WT and AD mice could be observed from the transaxial images. We conducted quantification with 3D ROIs and found that the total chemiluminescence signals from the 5xFAD mice group were 2.72-fold higher than that from the WT group at 10 minutes (Fig. 8b). We also quantified 2D images acquired at different wavelengths, and the largest difference was about 1.80-fold at 600 nm (SI Fig. S12). Our data suggested that 3D imaging not only can provide spatial information but also improve quantitative analysis.

**Figure 8.**
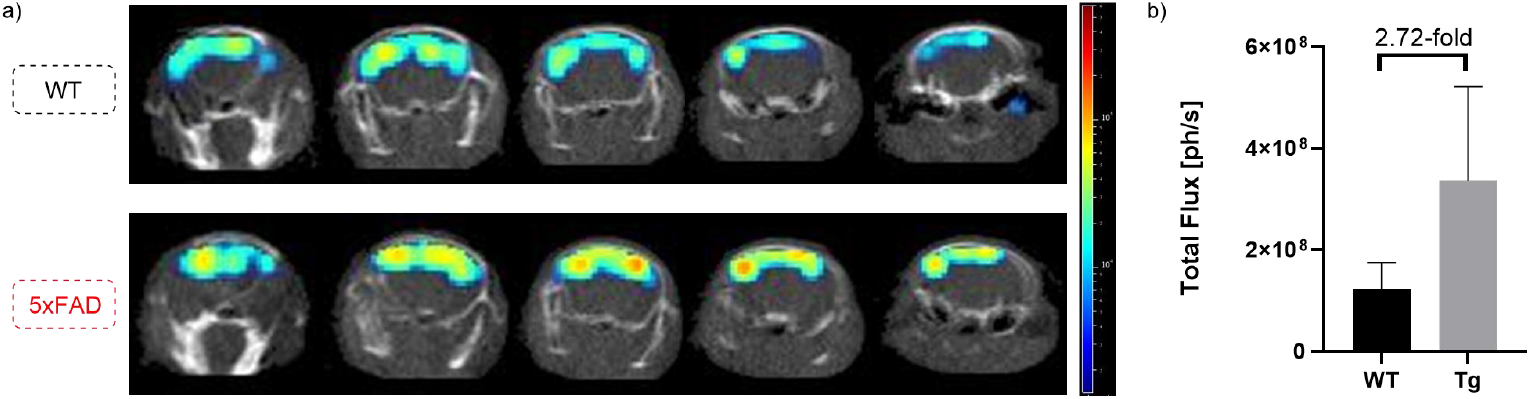
Difference between WT and AD mice via 3D imaging. a) Representative 3D brain image slices of transaxial view from a WT and a 5xFAD mice with ADLumin-5; b) Quantification with ROIs from the 3D reconstructed images of WT and 5xFAD mice. Data points with its error bar stand for mean ± s.d. derived from n = 3 independent animals.

## Discussion

Brain 3D tomography imaging at the macro- and meso-scale can be achieved with several imaging modalities, and each modality is often associated with certain limitations (1, 27). For example, PET imaging is very sensitive, but it is associated with high costs and radioactive materials (28). MRI is widely available, but few contrast agents are useful for molecular brain imaging due to the limited BBB penetration of the contrast agents (29). Photoacoustic imaging has recently been used for 3D brain tomography imaging (7); however, it has not been widely applied. The 3D brain tomography imaging in this report represents a new approach with certain advantages, including low-cost, fast, and excellent sensitivities. Remarkably, our results showed that the signals from the deep locations in the mice’s brains could be reconstructed, suggesting that it could be feasible to use our imaging probes and methods for larger animals such as rats and marmoset monkeys.

We believe that the success of 3D brain imaging is dependent on the high brightness of the probe. In our case, ADLumin-5’s in vivo radiance strength is about 157-fold higher than most of the reported chemiluminescence probes (including ADLumin-6 in this report). Although we have been successful in achieving 3D brain imaging in this report, several perspective improvements can be significantly beneficial for future studies. From our studies, we found that the signal intensity from each filter is critical for successful reconstructions. If the signal from each filter is lower than ∼ 1*10^5^ photon/s/cm^2^/sr, reliable reconstructions could be challenging. In our studies, ADLumin-6 showed the largest amplification upon incubating with Aβ aggregates; however, we experienced difficulties in our 3D brain imaging because ADLumin-6 was not bright enough to provide adequate photons for each filter. In this regard, further improving the QY and extending the emission wavelengths can enable more reliable 3D brain imaging. In addition, certainly, the 3D images in Fig. 7 and Fig. 8 still need further optimizations with better algorithms and better imaging systems to further improve the spatial resolutions. We believe that, with optimized probes and better reconstruction methods, it is possible to provide “PET-like” 3D brain optical imaging, which will have great potential for wide applications with low cost for preclinical brain disorder research.

## Materials and Methods

Materials, methods, and chemical synthesis are described in *SI Materials and Methods*. All animal experiments were approved by the Institutional Animal Use and Care Committee at Massachusetts General Hospital.

## Acknowledgments

This work was supported by NIH grants R01AG055413, R21AG059134 and R21AG078749 awards (to C. R.).

## Notes

### Competing Interest Statement

The authors have declared no competing interest.

