## Supplemental Information for "In Vivo Three-dimensional Brain Imaging with Chemiluminescence Probes in Alzheimer’s Disease Models"

**This PDF file includes:**

Supporting text
Figures S1 to S12
Tables S1
NMR spectra
SI References

**Materials and Methods**

Reagents used for the synthesis were purchased from Sigma-Aldrich and used without further purification. Column chromatography was performed on a glass column slurry-packed with silica gel (60 Å, 40–63 mm; SiliCycle Inc.). A$\beta_{1\text{-}40}$ TFA (A-1001-2) were purchased from rPeptide. Aβ aggregates for in vitro studies were generated by slowly stirring A$\beta_{40}$ in PBS buffer for 3 days at room temperature. $^1$H and $^{13}$C NMR spectra were recorded at 500 MHz and 125 MHz on Bruker spectrometers in $CDCl_3$, $CD_3OD$ or DMSO-$d_6$ solutions at room temperature with tetramethylsilane (TMS, $\delta$ = 0) as an internal standard. Liquid chromatography mass spectrometry (LC-MS) was performed using an Agilent 1200 Series apparatus with an LC/MSD trap and Daly conversion dynode detector with UV detection at 254 nm. Fluorescence measurements were carried out using an F-7100 fluorescence spectrophotometer (Hitachi). Transgenic female 5xFAD mice and age-matched wild-type female mice were purchased from Jackson Laboratory. All animal experiments were approved by the Institutional Animal Use and Care Committee at Massachusetts General Hospital. Animals were housed in group in standardized cages with a 12/12 h light/dark cycle with unrestricted access to food and water at room temperature (ca. 25°C) with ca. 55% humidity. The IVIS Spectrum animal imaging system (PerkinElmer) was used for in vitro and vivo imaging. 3D imaging was performed on an IVIS® Spectrum CT (PerkinElmer).

**Experimental section**

**Preparation of A$\beta_{40}$ aggregates.** 1.0 mg of A$\beta_{40}$ peptide (TFA) was suspended in a 1% ammonia hydroxyl solution (1.0 mL), then 100.0 µL of the resulting solution was taken and diluted 10-fold with PBS buffer (pH 7.4) and stirred at room temperature for 3 days. Transmission electron microscopy and fluorescence tests with Thioflavin T were used to confirm the formation of aggregates (1).

**In vitro fluorescence spectral test in solutions.** ADLumin-Xs/-Xs(O) solutions (250.0 nM) in 1.0 mL DMSO were prepared, and fluorescence spectra of the prepared solutions were recorded using an F-7100 fluorescence spectrophotometer (Hitachi). A blank control of DMSO was used to correct the final spectra.

**In vitro chemiluminescence spectral test in solutions.** ADLumin-Xs solutions (2.5 μM ) in 100 µL 10% DMSO/buffer with A$\beta_{40}$ aggregates (25.0 μM) were prepared, and chemiluminescence spectra of the prepared solutions were recorded with the IVIS SpectrumCT imaging system (PerkinElmer). A blank control of 10% DMSO/PBS was used to correct the final spectra.

**The measurement of pKa.** ADLumin-5/6 solutions (25.0 μM) in 3.0 mL 2.4% MeOH/pH buffer were prepared, and UV-vis spectra of the prepared solutions were recorded with UV-Vis spectrophotometer, and the spectra were scanned from 350 nm to 600 nm. The pKa was calculated through the fitting of the titration.

**Superoxide dismutases (SOD) inhibition test.** A solution of ADLumin-5 (2.5 μM, 10% DMSO) was incubated with A$\beta_{40}$ aggregates (25 μM), followed by adding superoxide dismutases (SOD) (0.5 μM) as $O_2^{-\bullet}$ inhibition. Then we measured the chemiluminescence intensity for each group. We found that the chemiluminescence intensity of the SOD group was lower than the group without SOD, which suggested that the chemiluminescence intensity was related to the concentration of $O_2^{-\bullet}$.

**Superoxide anion radical ($O_2^{-\bullet}$) detection by DHE.** Evaluation of $O_2^{-\bullet}$ was performed by the interaction between dihydroethidine and DNA (2). Dihydroethidium (DHE) was utilized as $O_2^{-\bullet}$ specific probe because it could intercalate in DNA and emit red fluorescence upon interacting with $O_2^{-\bullet}$. Briefly, ADLumin-5 (10 μM, 0.5% DMSO) and DHE (25.0 μM, 0.05% DMSO) were dissolved in PBS solutions (2.0 mL) containing ctDNA (250.0 μg/mL). A mixture containing DHE and ctDNA only was used as the control. The fluorescence spectra were measured every 30 seconds.

**Singlet oxygen ($^1O_2$) detection by SOSG.** $^1O_2$ was tested using SOSG as the fluorescent indicator. The solution of ADLumin-5 (2.0 μM, 0.5% DMSO) in PBS solutions (2.0 mL) containing SOSG (10.0 μM, 0.05% DMSO) was added to the cuvette. A solution containing SOSG (10.0 μM) only was used as the control. The fluorescence spectra were measured every 15 seconds.

**Quantum yields detection.** Chemiluminescence quantum yields were determined by using luminol with $H_2O_2$ as oxidant and hemin as a catalyst as the standard with a known QY of 1.14 × $10^{-2}$ einsteins/mol in PBS buffers (pH 11.6). All solutions: luminol (0.025 mM in pH 11.6 PBS buffer), hemin solutions with absorbances 0.6 at 414 nm, and $H_2O_2$ (10.0 mM). The 0.6 mL of the luminol solution was placed in the luminometer in the dark. $H_2O_2$ solution (100.0 μL) was rapidly added to the luminol solution, followed by hemin solution (100.0 μL). A burst of light appeared and decayed relatively quickly. An average of at least ten measurements were used to determine the luminol calibration factor (flum) according to the following equations. Then, chemiluminescence intensities of ADLumin-X (20.0 μM, 10% DMSO) were measured in the presence of $A\beta_{40}$ (25.0 μM) at 37 °C, which were plotted as a function of time, respectively. Then, according to the first equation, our probe's chemiluminescence quantum yields can be calculated.

$$\Phi_{cL} = \frac{Q \times f_{lum} \times f_{photo}}{n} \text{ (einsteins/mol)}$$

$$f_{lum} = \frac{\Phi_{lum} \times n_{lum}}{Q_{lum}}$$

$\Phi_{CL}$: the chemiluminescence QYs,

Q: the total light emission obtained by integration of emission intensity.

$f_{lum}$ is obtained by measuring the emission kinetics of the luminol reaction performed in standard conditions.

$f_{photo}$ is given by the manufacturer.

n: the number of moles of luminol ($n_{lum}$).

**Mimic tissue penetration studies.** A tube that had similar intensity of emitted light from ADLumin-5 or -6 was placed under the abdomen of a nude mouse (the depth is about 1.5 cm), and then the signals were collected with the IVIS®SpectrumCT imaging system, which captured the signals from the dorsal side.

**In vitro histological staining.** A 25-μm brain slice from an 18-month-old APP/PS1 mouse was fixed with 4% formalin for 5 min and washed with distilled water twice for 5 min. The slice was incubated with a CRANAD-2 solution (2.5 μM in 10% DMSO and 90% PBS) for 15 min and then, washed with 40% ethanol, followed by washing with double distilled (dd) water. After drying, the same slide was incubated with ADLumin-5/-6 (250 nM in 10% DMSO and 90% PBS) for 10 min. After drying, the slice was covered with FluoroShield mounting medium (Abcam) and sealed with nail polish. Fluorescence images were obtained using the Olympus IX83 Microscope System with red and green channels.

**In vivo 2D mouse studies.** In vivo chemiluminescence imaging was performed using the IVIS®SpectrumCT animal imaging system. Balb/c mice (8-month-old, female) and age-matched 5xFAD mice were i.p. injected ADLumin-5/-6 (4.0 mg/kg, 5% DMSO, and 95% PBS), respectively. Then background images were acquired with an open filter. Chemiluminescence imaging was performed at 5, 10, 20, 30, 60, 90, and 180 min after the i.p. injection of ADLumin-5/-6 (4.0 mg/kg, 5% DMSO, and 95% PBS). Living Image 4.7.1 software (PerkinElmer) was used for the data analysis.

**In vivo 3D mouse imaging studies.** In vivo chemiluminescence imaging was performed using the IVIS® SpectrumCT. Balb/c mice and 5xFAD mice (8-month-old, female) were i.v. injected ADLumin-5 (4.0 mg/kg, 5% DMSO, and 95% PBS). Chemiluminescence signals were collected with 540-, 560-, 580-, 600-, 620-, 640-, 660-, 680- and 700-nm filters, and a signal threshold < 10.0% was used for the 3D reconstruction. Chemiluminescence imaging was performed 10 min after the i.v. injection of ADLumin-5 (4.0 mg/kg, 5% DMSO, and 95% PBS). The acquisition time was 60 seconds for each filter, and the radiance from each filter was > $1*10^5$ photon/s/$cm^2$/sr. Upon completing the image collection, 3D reconstruction was performed with the diffuse luminescence tomographic algorithm (DLIT™) algorithm.

**Density functional theory (DFT) calculations.** Density functional theory (DFT) and time-dependent density functional theory (TD-DFT) calculations were performed using the Q-Chem Software package (3). All calculations were performed using the conductor-like polarizable continuum model (C-PCM) (4) with the dielectric constant of DMSO (46.7); in addition, the optical dielectric constant of DMSO (3.9) was used for the absorption (i.e., vertical excitation) energy calculations. For each molecule, the ground-state geometry was optimized using DFT with the ωB97X-D functional (5) and 6-31+G* basis set (6). At the optimized geometries, TD-DFT calculations were performed using the same functional and basis set to obtain vertical excitation energies. For each molecule, the lowest-energy excited state with a significant oscillator strength (greater than 0.7) was chosen to analyze absorption peaks and as the initial state for subsequent emission wavelength calculations. For the emission spectra calculations, we started from the ground-state geometries and performed further geometry optimization on the chosen excited state potential energy surface. In order to ensure the optimization proceeded on the same surface, we used the state tracking algorithm implemented in Q-Chem, which is based on the transition density overlap in the atomic orbital (AO) representation between consecutive geometries.

$$S_{12}^{ij} = Tr\left(R_1^i\, S_{12}\, R_2^{j\top}\, S_{21}\right)$$

Here, $S_{12}$ and $S_{21}$ are overlap matrices between basis functions at the two geometries, $R_1^i$ is the transition density matrix for excitation to state *i* at the previous geometry, and $R_2^{j\top}$ is the transposed transition density matrix for excitation to state *j* at the current geometry. The orbital plots were generated using the IQmol graphical user interface. The HOMO-LUMO energy gaps shown in Fig. 2c were calculated at the optimized excited state geometries.

**Substrate Preparations and Characterizations.**
**The synthetic route**

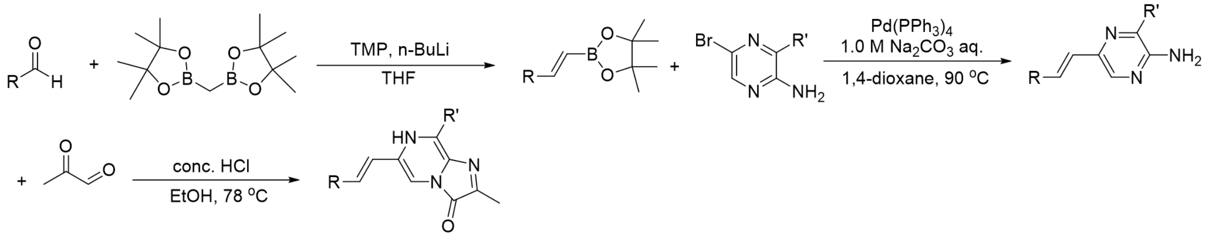


**General method for preparation of ADLumin-X.**
**Step I**

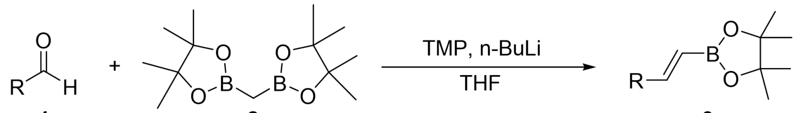


1 2 3

To a solution of TMP in anhydrous THF (35 mL), n-BuLi was added at 0 °C. The resulting mixture was stirred for 5 mins at 0 °C, followed by an addition of a solution of bis-(4,4,5,5-tetramethyl-1,3,2-dioxaborolan-2-yl)methane (10.0 mmol, 1.2 equiv.) in THF (20.0 mL). The resulting solution was stirred at 0 °C for 15 minutes. Then the reaction vial was cooled to -78 °C, and a solution of aldehyde **1** (8.3 mmol, 1.0 equiv.) in THF (10 mL) was added. The reaction vial was stirred at -78°C for additional 4 hours. Upon completion, the reaction mixture was concentrated under reduced pressure, and the 1,2-disubtituted-vinyl boronate products **3** were purified by flash silica chromatography.

**Step II**

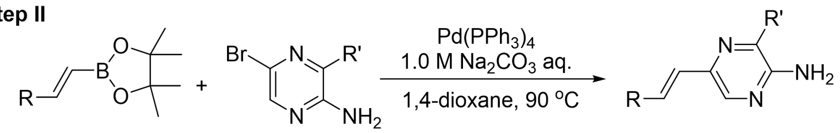


**3** **4** **5**

Under an argon atmosphere, compound **3** (4.4 mmol, 2.0 equiv.) and compound **4** (2.2 mmol, 1.0 equiv.) were dissolved in a mixture of 1,4-dioxane (20.0 mL), and 1.0 M $Na_2CO_3$ aqueous solution (6.6 mL), and the resulting mixture was degassed in vacuo. A catalytic amount of tetrakis(triphenylphosphine)palladium(0) (10% equiv.) was added to the mixture, and the mixture was heated at 90 °C for 6 h. After cooling, the mixture was diluted with EtOAc (30 mL x 3) and washed with water and brine, dried over $Na_2SO_4$, and evaporated. The resulting residue was purified by flash silica chromatography to obtain product **5**.

**Step III**

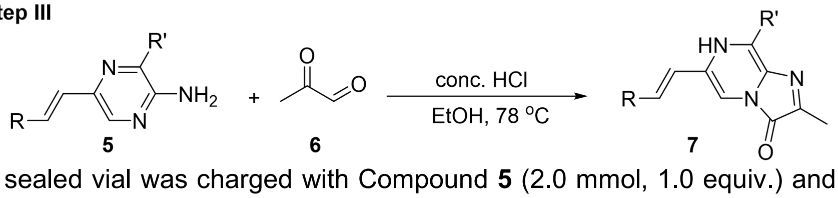


**5** **6** **7**

A sealed vial was charged with Compound **5** (2.0 mmol, 1.0 equiv.) and **6** (6.0 mmol, 3.0 equiv.) in EtOH (15 mL) which was bubbled with $N_2$, and then the reaction mixture was stirred at room temperature for 10 min under an argon atmosphere. Adding concentrated HCl (7.2 mmol, 3.6 equiv.) in EtOH (6 mL) to the mixture via syringe over 10 minutes. The mixture was refluxed at 78 °C 4-12 h. LC-MS monitored the reaction. Upon completion, the mixture was cooled to room temperature and concentrated the crude under vacuum. The desired product **7** was precipitated by adding $Et_2O$ to the residue dissolved in a minimum of MeOH.

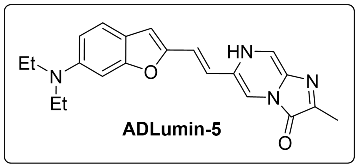

**ADLumin-5**

**Step I**

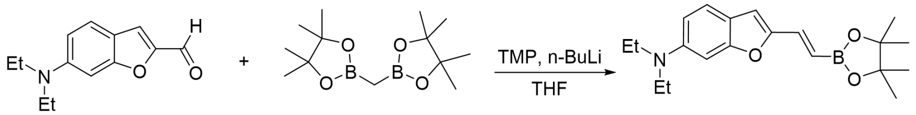


**8** **9**

**(*E*)-N,N-diethyl-2-(2-(4,4,5,5-tetramethyl-1,3,2-dioxaborolan-2-yl)vinyl)benzofuran-6- amine (9)**. Following the general method in step (**I**), the reaction of Compound **8** (7) (1.81 g, 8.3 mmol) afforded product **9** as a light yellow solid (1.49 g, 52% yield): TLC $R_f$ = 0.60 (EtOAc/hexanes = 1/4); $^1$H NMR (500 MHz, $CDCl_3$) δ 7.32 (d, *J* = 8.7 Hz, 1H), 7.19 (d, *J* = 18.1 Hz, 1H), 6.72 (d, *J* = 1.5 Hz, 1H), 6.65 (dd, *J* = 8.7, 2.2 Hz, 1H), 6.57 (s, 1H), 6.09 (d, *J* = 18.1 Hz, 1H), 3.40 (q, *J* = 7.1 Hz, 4H), 1.31 (s, 12H), 1.19 (t, *J* = 7.1 Hz, 6H); $^{13}$C NMR (125 MHz, $CDCl_3$) 158.0, 153.0, 147.4, 136.9, 121.7, 121.7, 118.2, 109.8, 108.0, 93.9, 83.4, 45.1, 25.0, 12.7.

**Step II**

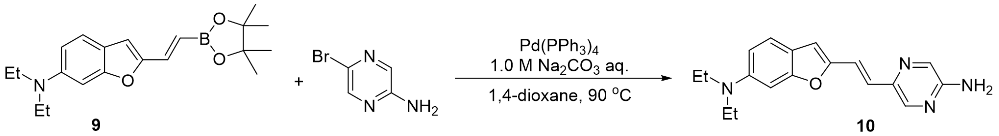


**9** **10**

**(*E*)-5-(2-(6-(diethylamino)benzofuran-2-yl)vinyl)pyrazin-2-amine (10)**. Following the general method in step (**II**), the reaction of Compound **9** (1.49 g, 4.4 mmol) afforded product **10** as a brown solid (0.61 g, 90% yield): TLC $R_f$ = 0.31 (EtOAc/hexanes = 1/1); $^1$H NMR (500 MHz, $CDCl_3$) δ 8.00 (d, *J* = 6.1 Hz, 2H), 7.36 – 7.28 (m, 2H), 7.07 (d, *J* = 15.7 Hz, 1H), 6.75 (s, 1H), 6.65 (d, *J* = 8.1 Hz, 1H), 6.57 (s, 1H), 4.60 (br, 2H, -$NH_2$), 3.41 (q, *J* = 7.0 Hz, 4H), 1.20 (t, *J* = 7.0 Hz, 6H);

$^{13}C$ NMR (125 MHz, $CDCl_3$) 157.6, 152.9, 152.7, 147.0, 141.4, 132.4, 122.5, 121.2, 118.6, 117.8, 109.7, 106.8, 93.9, 45.1, 29.8, 12.7.

**Step III**

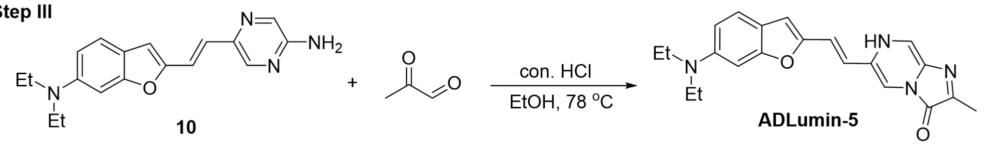


**(*E*)-6-(2-(6-(diethylamino)benzofuran-2-yl)vinyl)-2-methylimidazo[1,2-a]pyrazin-3(7H)-one (ADLumin-5).** Following the general method in step (III), the reaction of Compound **10** (0.61 g, 2.0 mmol) afforded **ADLumin-5** as an orange-brown solid (0.16 g, 22% yield): $^1H$ NMR (500 MHz, $CD_3OD$) $\delta$ 9.04 (s, 1H), 8.54 (s, 1H), 7.93 (s, 1H), 7.88 (d, *J* = 8.4 Hz, 1H), 7.74 (d, *J* = 15.6 Hz, 1H), 7.52 (d, *J* = 15.6 Hz, 2H), 7.14 (s, 1H), 3.76 (m, 4H), 2.54 (s, 3H), 1.19 (t, *J* = 7.2 Hz, 6H); $^{13}C$ NMR (125 MHz, $CD_3OD$) 158.4, 156.2, 135.5, 132.3, 128.5, 125.9, 124.3, 121.7, 118.6, 114.9, 108.3, 107.1, 55.3, 10.8, 9.8; HRMS-ESI (m/z) [$M+H^+$]: calcd for $C_{21}H_{23}N_4O_2$ 363.1816, found 363.1813.

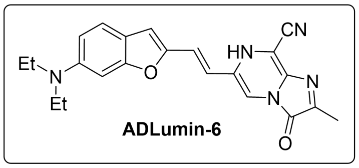


**Step I**

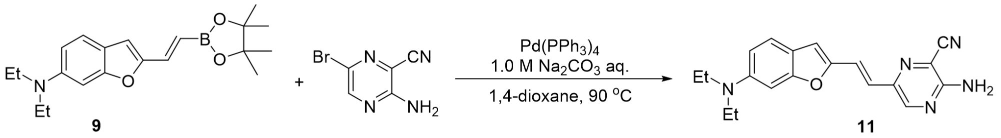


**(*E*)-3-amino-6-(2-(6-(diethylamino)benzofuran-2-yl)vinyl)pyrazine-2-carbonitrile (11).** Following the general method in step (II), the reaction of Compound **9** (1.50 g, 4.4 mmol) afforded product **11** as an orange-red solid (0.57 g, 78% yield): TLC $R_f$ = 0.49 (EtOAc/hexanes = 1/1); $^1H$ NMR (500 MHz, $CDCl_3$) $\delta$ 8.23 (s, 1H), 7.40 – 7.32 (m, 2H), 7.02 (d, *J* = 15.6 Hz, 1H), 6.73 (s, 1H), 6.67 (d, *J* = 8.8 Hz, 2H), 3.42 (q, *J* = 7.0 Hz, 4H), 1.21 (t, *J* = 7.0 Hz, 6H); $^{13}C$ NMR (125 MHz, $CDCl_3$) 157.9, 154.4, 151.9, 147.5, 145.8, 143.2, 121.6, 120.5, 120.1, 118.4, 115.5, 113.1, 109.9, 108.7, 93.5, 45.1, 12.7.

**Step II**

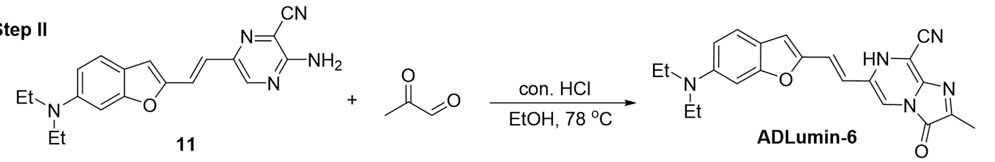


**(*E*)-6-(2-(6-(diethylamino)benzofuran-2-yl)vinyl)-2-methyl-3-oxo-3,7-dihydroimidazo[1,2-a]pyrazine-8-carbonitrile (ADLumin-6).** Following the general method in step (II), the reaction of Compound **11** (0.57 g, 1.7 mmol) afforded **ADLumin-6** as a brown-red solid (0.10 g, 15% yield): $^1H$ NMR (500 MHz, $CD_3OD$) $\delta$ 8.58 (s, 1H), 7.86 (d, *J* = 8.6 Hz, 2H), 7.72 (d, *J* = 15.6 Hz, 1H), 7.49 (d, *J* = 15.6 Hz, 1H), 7.45 (dd, *J* = 8.6, 1.9 Hz, 1H), 7.12 (s, 1H), 3.75 (m, 4H), 2.52 (s, 3H), 1.17 (d, *J* = 7.2 Hz, 6H); $^{13}C$ NMR (125 MHz, $CD_3OD$) 158.4, 156.2, 140.2, 139.9, 135.5, 132.3, 129.8, 126.1, 124.3, 122.3, 122.0, 118.6, 117.6, 113.7, 108.5, 107.1, 55.3, 10.8, 10.2; HRMS-ESI (m/z) [$M+H^+$]: calcd for $C_{22}H_{22}N_5O_2$ 388.1768, found 388.1776.

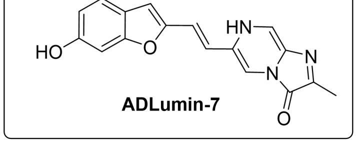

**Step I**

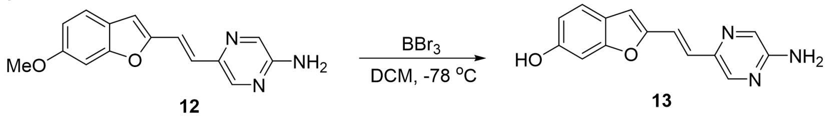

12 13

**(*E*)-2-(2-(5-aminopyrazin-2-yl)vinyl)benzofuran-6-ol (13)**. A $BBr_3$ solution (1.0M, 5.7 mmol, 10.0 equiv.) was added to Compound **12** (0.15g, 0.6 mmol) in the form of $CH_2Cl_2$ solution at -78 °C, followed by stirring at room temperature for 3 hours. Then the residue was extracted with EtOAc (15 mL x 3) and the yellow phase was washed with water and brine, dried over $Na_2SO_4$ and evaporated. The resulting yellow product **13** was directly subjected to the next reaction without further purification.

**Step II**

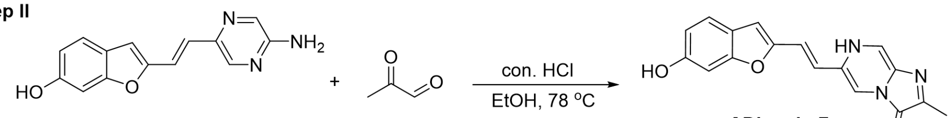

13 ADLumin-7

**(*E*)-6-(2-(6-hydroxybenzofuran-2-yl)vinyl)-2-methylimidazo[1,2-a]pyrazin-3(7H)-one (ADLumin-7)**. Following the general method in step (II), the reaction of Compound **13** (0.14 g, 0.6 mmol) afforded **ADLumin-7** as a brown-red solid (0.13 g, 74% yield for two steps): $^1H$ NMR (500 MHz, $CD_3OD$) δ 8.94 (s, 1H), 8.38 (s, 1H), 7.57 (d, *J* = 15.7 Hz, 1H), 7.38 (d, *J* = 8.4 Hz, 1H), 7.23 (d, *J* = 15.7 Hz, 1H), 6.88 (d, *J* = 1.8 Hz, 1H), 6.86 (s, 1H), 6.76 (dd, *J* = 8.4, 1.8 Hz, 1H), 2.53 (s, 3H); $^{13}C$ NMR (125 MHz, $CD_3OD$) 159.4, 159.1, 155.2, 140.9, 140.5, 136.9, 129.4, 123.9, 123.8, 123.8, 122.5, 114.9, 114.9, 110.8, 110.1, 99.4, 11.2; HRMS-ESI (m/z) [M+H$^+$]: calcd for $C_{17}H_{14}N_3O_2$ 308.1030, found 308.1029.

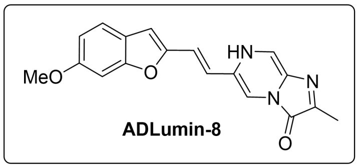

**Step I**

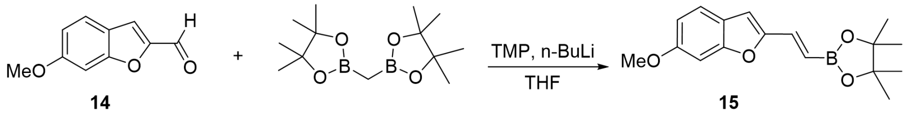

14 15

**(*E*)-2-(2-(6-methoxybenzofuran-2-yl)vinyl)-4,4,5,5-tetramethyl-1,3,2-dioxaborolane (15)**. Following the general method in step (I), the reaction of Compound **14** (8) (0.96 g, 5.4 mmol) afforded product **15** as a white solid (1.54 g, 95% yield): TLC $R_f$ = 0.45 (EtOAc/hexanes = 1/4); $^1H$ NMR (500 MHz, $CDCl_3$) δ 7.39 (d, *J* = 8.6 Hz, 1H), 7.22 (d, *J* = 18.2 Hz, 1H), 6.99 (s, 1H), 6.84 (dd, *J* = 8.6, 2.2 Hz, 1H), 6.64 (s, 1H), 6.20 (d, *J* = 18.2 Hz, 1H), 3.86 (s, 3H), 1.31 (s, 12H); $^{13}C$ NMR (125 MHz, $CDCl_3$) 159.0, 156.5, 154.6, 136.6, 122.2, 121.7, 112.4, 107.4, 95.8, 83.6, 55.8, 24.9.

**Step II**

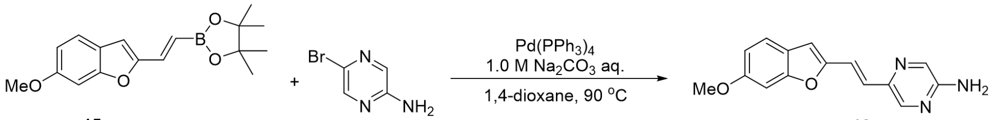

15 12

**(*E*)-5-(2-(6-methoxybenzofuran-2-yl)vinyl)pyrazin-2-amine (12)**. Following the general method in step (II), the reaction of Compound **15** (1.54 g, 3.9 mmol) afforded product **12** as a yellow solid (0.32 g, 62% yield): TLC $R_f$ = 0.19 (EtOAc/hexanes = 1/1); $^1H$ NMR (500 MHz, $CDCl_3$) δ 8.03 (s, 2H), 7.39 (d, *J* = 8.5 Hz, 1H), 7.34 (d, *J* = 15.6 Hz, 1H), 7.14 (d, *J* = 15.6 Hz, 1H), 7.01 (d, *J* = 2.1 Hz, 1H), 6.84 (dd, *J* = 8.5, 2.1 Hz, 1H), 6.64 (s, 1H), 4.71 (br, 2H, $-NH_2$), 3.87 (s, 3H); $^{13}C$ NMR

(125 MHz, $CDCl_3$) 158.7, 156.3, 154.4, 152.9, 141.1, 132.7, 124.1, 122.7, 121.2, 117.7, 112.0, 106.4, 95.8, 55.9.

**Step III**

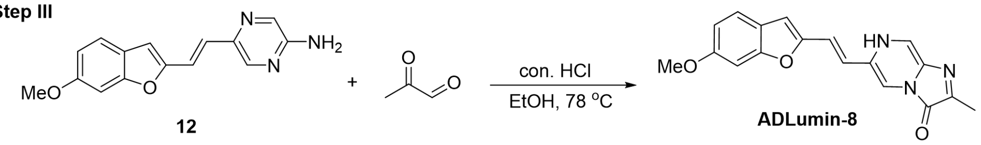


**12** **ADLumin-8**

(E) **-6-(2-(6-methoxybenzofuran-2-yl)vinyl)-2-methylimidazo[1,2-a]pyrazin-3(7H)-one (ADLumin-8).** Following the general method in step (**III**), the reaction of Compound **12** (0.16 g, 0.6 mmol) afforded **ADLumin-8** as a brown solid (0.12 g, 63% yield): $^1H$ NMR (500 MHz, $CD_3OD$) $\delta$ 8.88 (s, 1H), 8.30 (s, 1H), 7.53 (d, *J* = 15.7 Hz, 1H), 7.40 (d, *J* = 8.6 Hz, 1H), 7.18 (d, *J* = 15.7 Hz, 1H), 7.01 (s, 1H), 6.84 (s, 1H), 6.82 (dd, *J* = 8.6, 1.9 Hz, 1H), 3.85 (s, 3H), 2.49 (s, 3H); $^{13}C$ NMR (125 MHz, $CD_3OD$) 160.7, 157.8, 154.4, 135.0, 128.4, 125.2, 123.4, 122.6, 122.5, 121.5, 113.6, 113.4, 109.3, 96.3, 56.2, 10.2; HRMS-ESI (m/z) [M+H$^+$]: calcd for $C_{18}H_{16}N_3O_3$ 322.1186, found 322.1181.

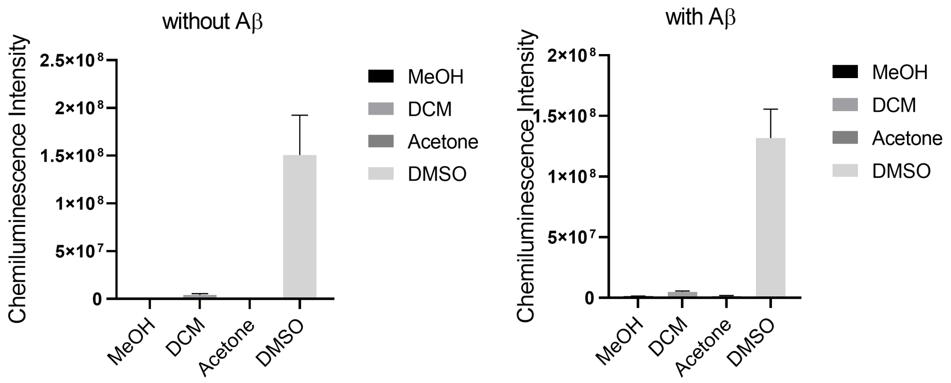


**Fig. S1.** Chemiluminescence of ADLumin-5 (2.5 μM, 10% DMSO) incubated with/without $A\beta_{40}$ in PBS buffer with different co-solvents. Data are represented as mean ± s. d. with n = 3 independent samples.

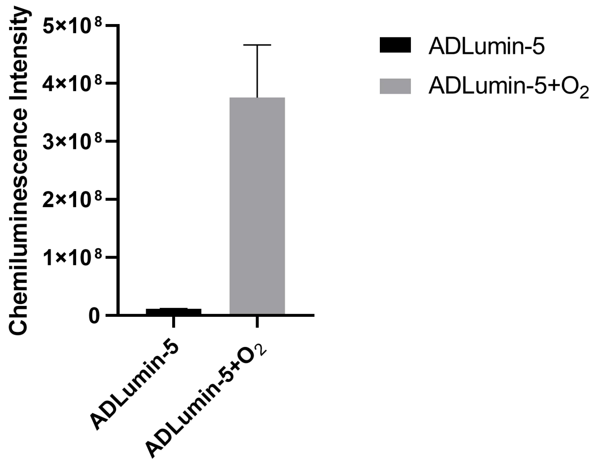


**Fig. S2.** The chemiluminescence of ADLumin-5 with and without $O_2$ bubbling.

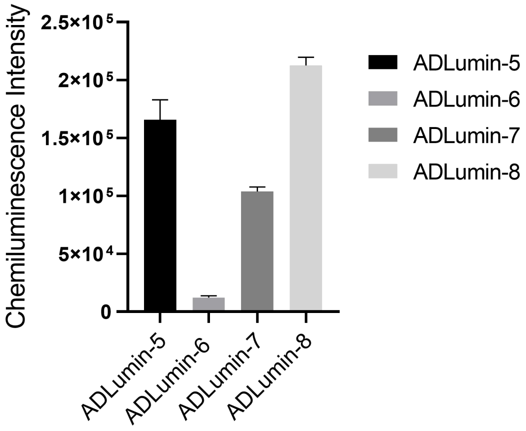


**Fig. S3.** Chemiluminescence intensities of ADLumin-Xs in PBS buffers.

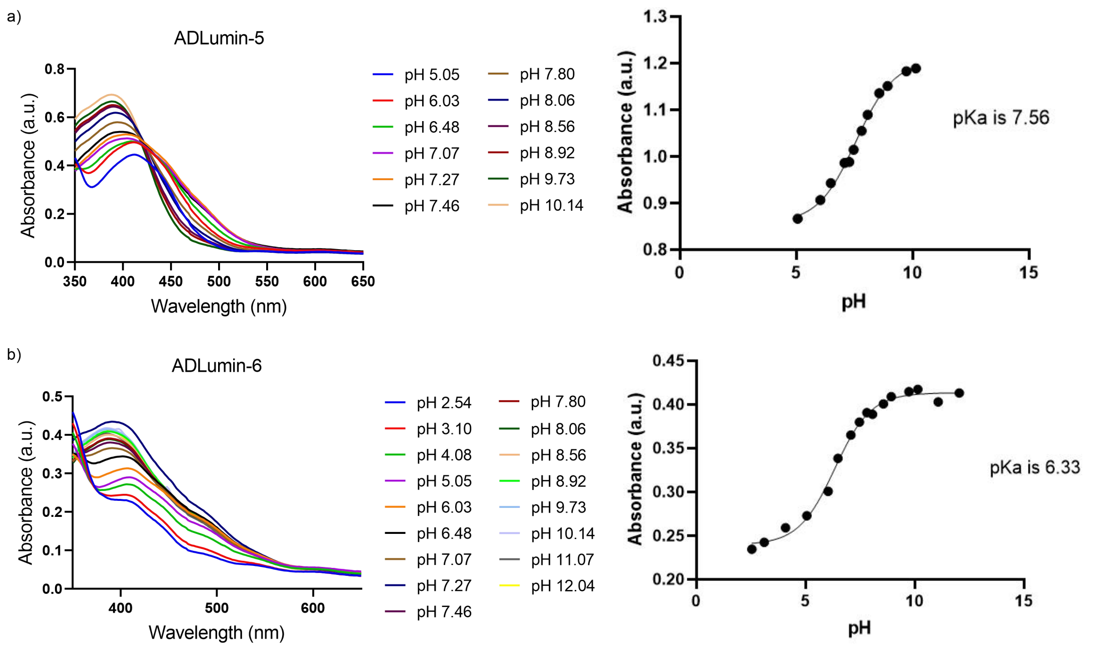


**Fig. S4.** The measurement of pKa of ADLumin-5 and ADLumin-6. a) Absorbance spectra of ADLumin-5 under different pH (left panel) and the fitting curve for pKa calculation (right panel); b) Absorbance spectra of ADLumin-6 under different pH (left panel) and the fitting curve for pKa calculation (right panel).

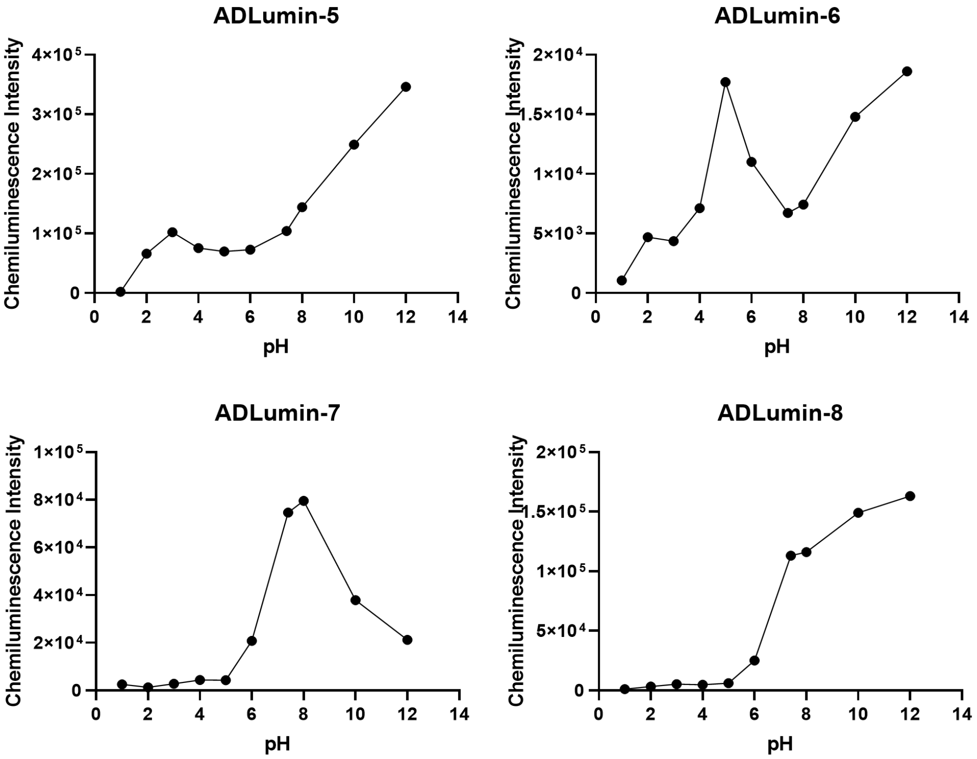


**Fig. S5.** The pH dependence of chemiluminescence intensities of ADLumin-Xs (X = -5, -6, -7, and -8).

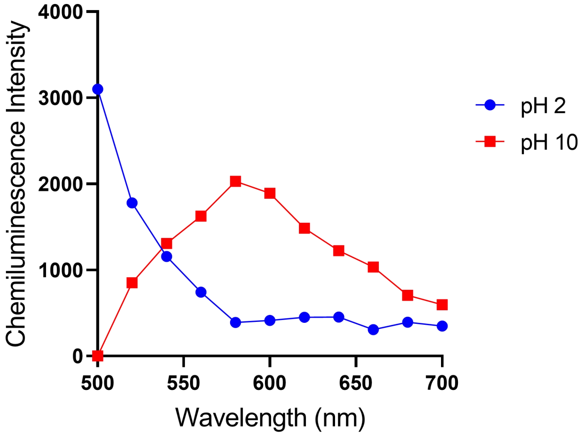


**Fig. S6.** The chemiluminescence emission spectra of ADLumin-5 in PBS buffers of pH 2 and 10, respectively.

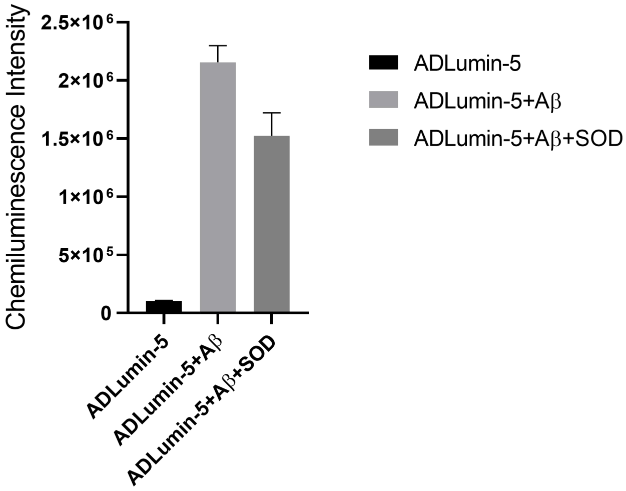


**Fig. S7.** The chemiluminescence intensity of ADLumin-5 in the presence of $A\beta_{40}$ aggregates and superoxide dismutase (SOD), which was used to scavenge superoxide radicals ($O_2^{-\bullet}$).

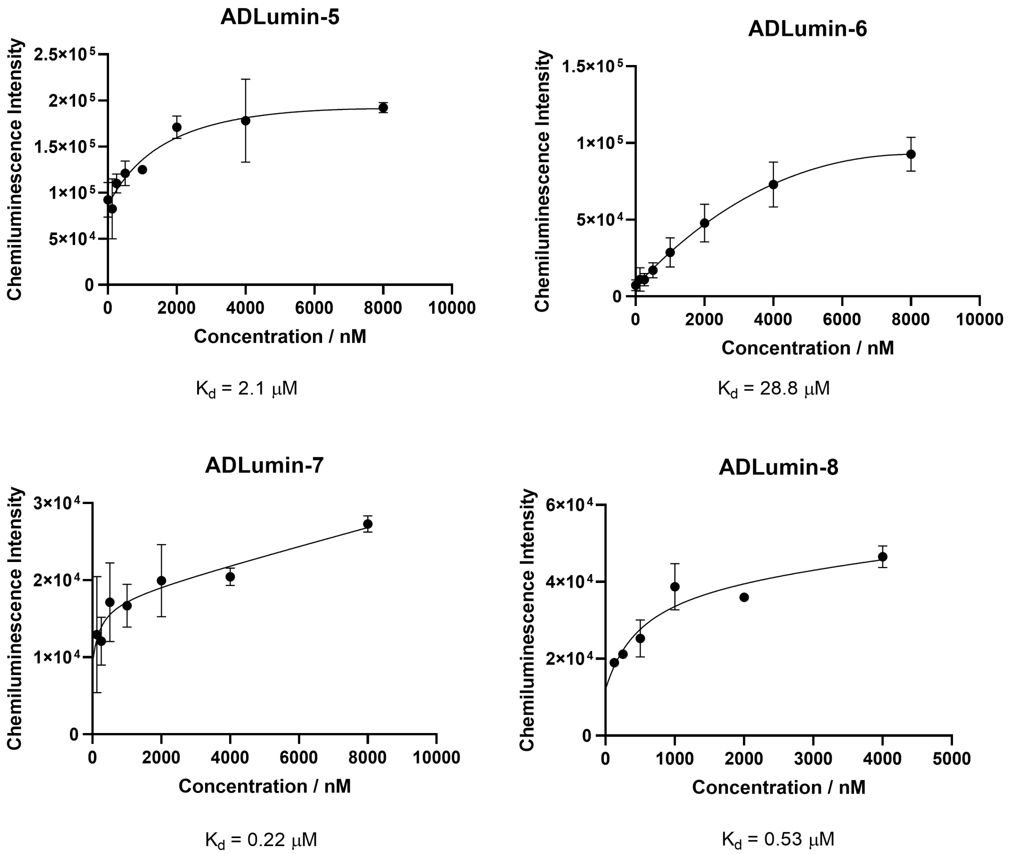


**Fig. S8.** Binding constant Kd measurement of ADLumin-X in the presence of Aβ aggregates. Binding affinity assays of ADlumin-X with $A\beta_{40}$ aggregates were performed with the chemiluminescence intensities of ADLumin-X (250 nM) with increasing concentrations of $A\beta_{40}$ aggregates from 0 to 8 μM. The data point with its error bar indicates mean ± s.d. derived from n=3 biologically independent samples.

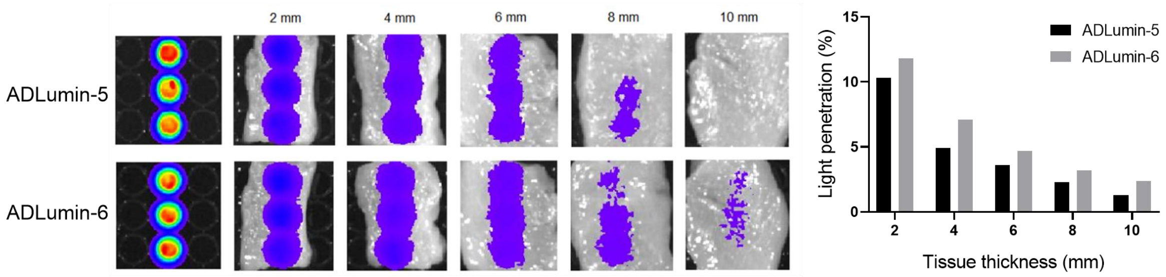


**Fig. S9.** The data point with its error bar indicates mean ± s.d. derived from n=3 biologically independent samples. The measurement of chemiluminescence penetration of ADLumin-5 (upper panel) and ADLumin-6 (lower panel) with chicken breast tissues through in vitro imaging (left panel) and the corresponding quantification (right panel).

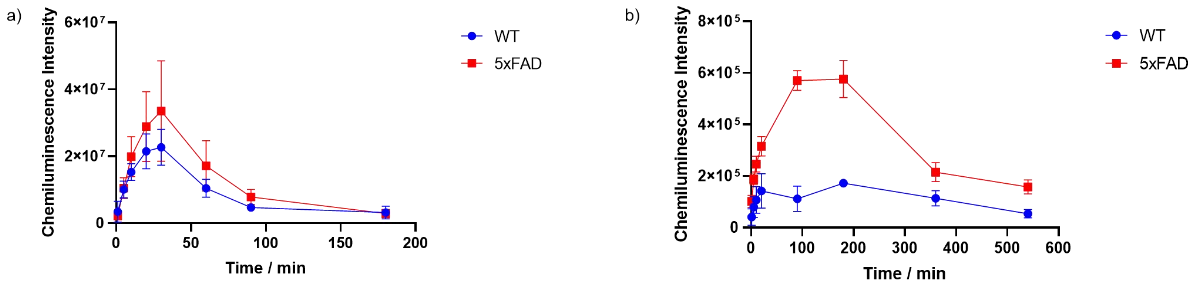


**Fig. S10.** a) The time course of ADLumin-5' chemiluminescence signals through in vivo imaging with 5xFAD and WT mice after i.p. injection; b) The time course of ADLumin-6' chemiluminescence signals through in vivo imaging with 5xFAD and WT mice after i.p. injection.

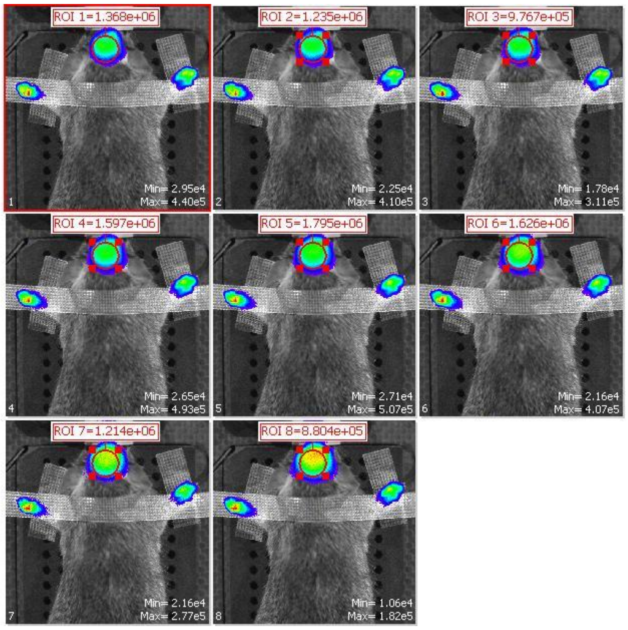


**Fig. S11.** The raw 2D images of each emission filter with different wavelengths of a 5xFAD mouse using ADLumin-5.

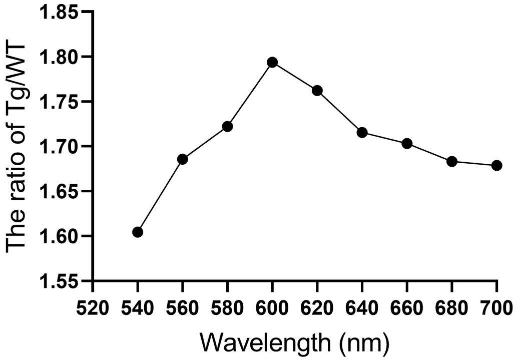


**Fig. S12.** The quantification of the chemiluminescence signal ratio of ADLumin-5 between 5xFAD and WT by raw 2D images of each emission filter with different wavelengths.

**Table S1.** Computed emission energies, wavelengths, oscillator strengths, and amplitudes for each contributing electronic transition for ADLumin-X(O) emissions in DMSO with 6-31+G* basis set.

| | ωB97X-D | | | | |
|---|---|---|---|---|---|
| Product | $\Delta E$ (eV) | $\lambda$ (nm) | f | O→V (amp.) | $\Delta E_{O \to V}$ (eV) |
| 5(O) | 2.231 | 555.7 | 1.831 | H→L (0.9641) | 5.434 |
| 6(O) | 2.116 | 585.9 | 1.787 | H→L (0.9616) | 5.309 |
| 7(O) | 2.502 | 495.5 | 1.832 | H→L (0.9757) | 5.845 |
| 8(O) | 2.487 | 498.6 | 1.874 | H→L (0.9742) | 5.815 |

### NMR spectra of ADLumin-X

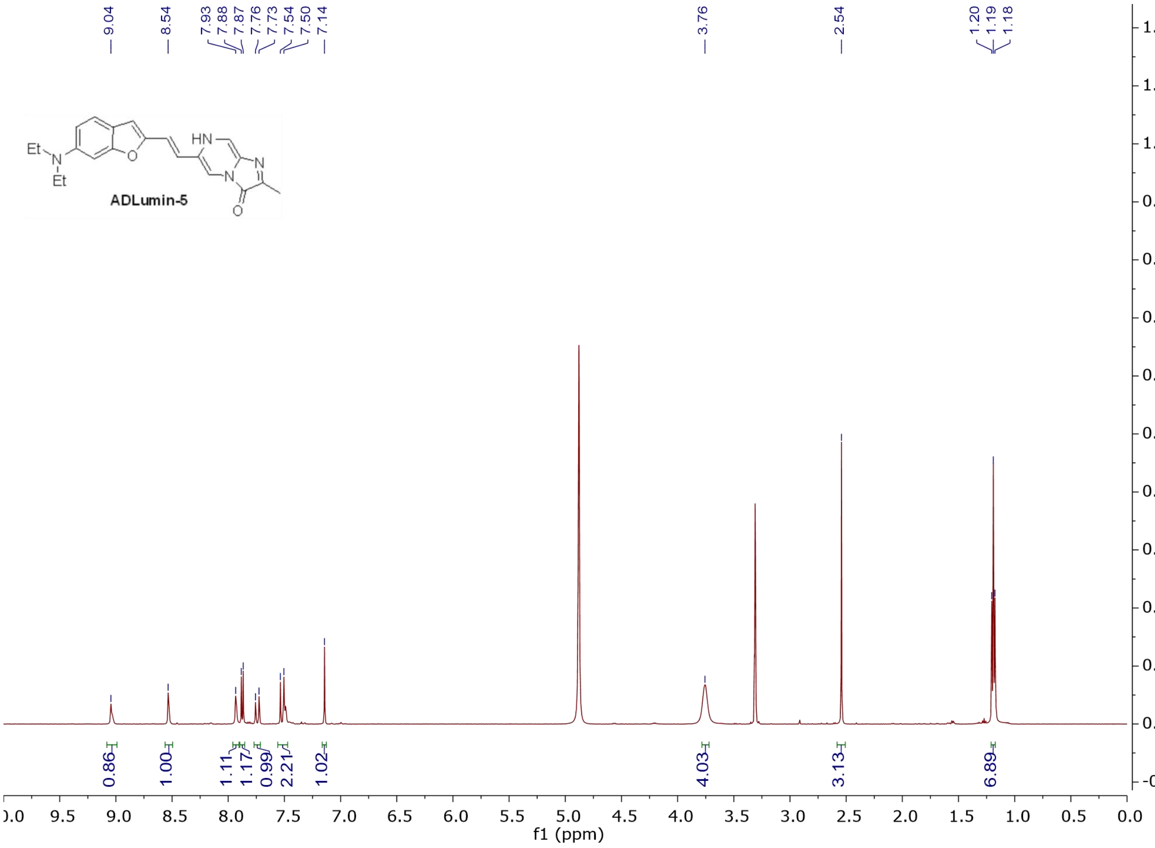

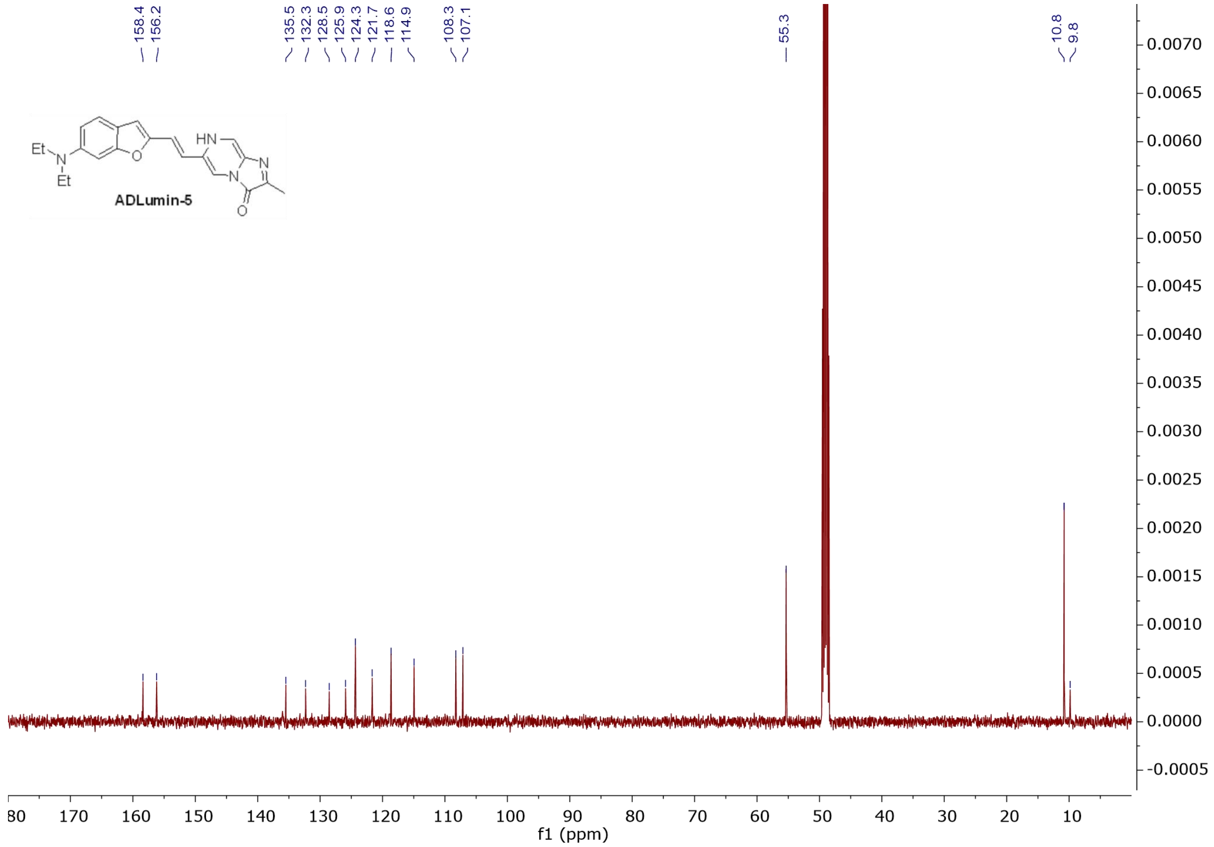
158.4
156.2
135.5
132.3
128.5
125.9
124.3
121.7
118.6
114.9
108.3
107.1
55.3
10.8
9.8
Et
N
Et
O
HN
N
N
O
ADLumin-5
f1 (ppm)

Spectrum

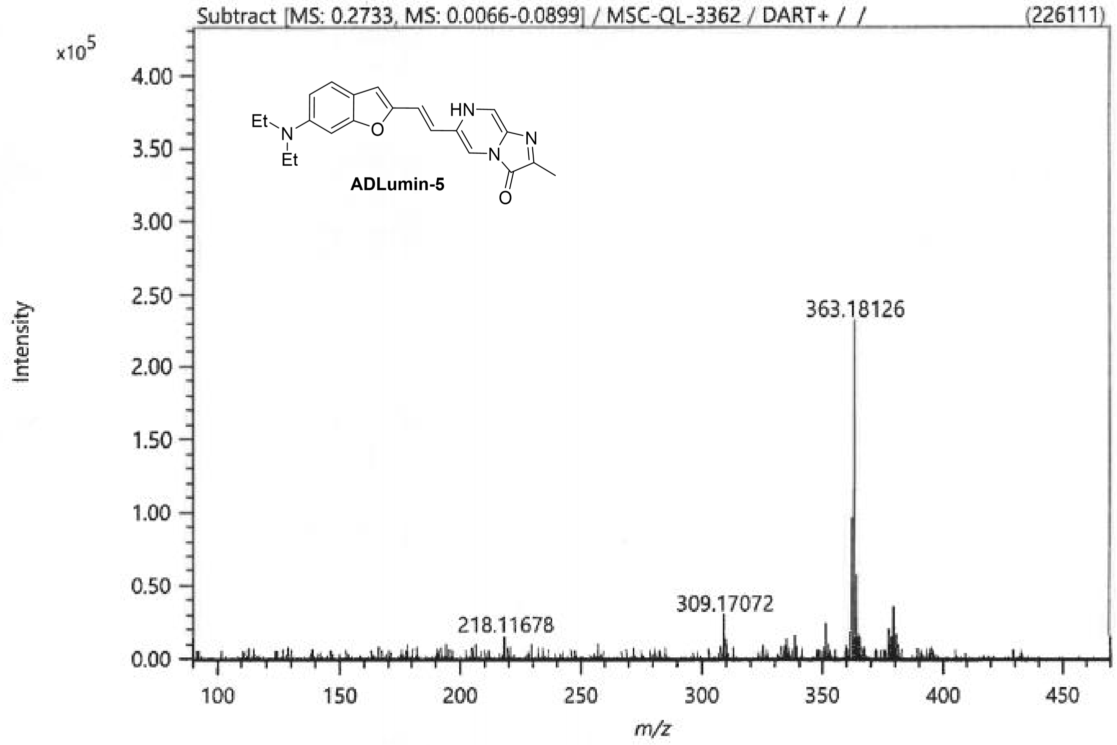


Elemental Composition

| Parameters | |
|---|---|
| Tolerance: | 6.00 ppm |
| Electron: | Odd/Even |
| Charge: | +1 |
| DBE: | -50.5 - 150.0 |

Elements Set 1:

| Symbol | C | H | O | N | S | Br |
|---|---|---|---|---|---|---|
| Min | 1 | 1 | 1 | 1 | 0 | 0 |
| Max | 100 | 100 | 2 | 5 | 0 | 0 |

Results

| Mass | Intensity | Formula | Calculated Mass | Mass Difference [ppm] | DBE |
|---|---|---|---|---|---|
| 363.18126 | 226110.82 | C21 H23 N4 O2 | 363.18155 | -0.81 | 12.5 |

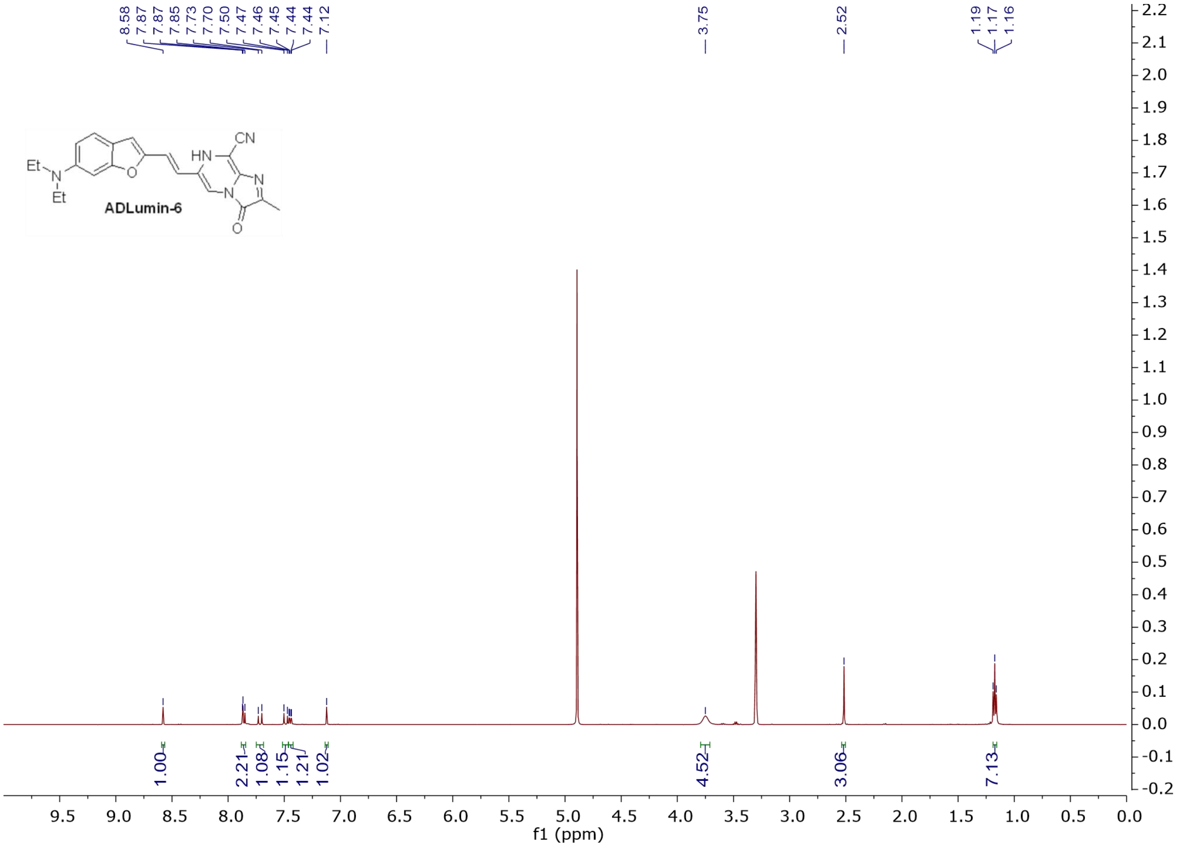

8.58
7.87
7.87
7.85
7.73
7.70
7.50
7.47
7.46
7.45
7.44
7.44
7.12
3.75
2.52
1.19
1.17
1.16
CN
HN
N
N
O
O
Et
N
Et
ADLumin-6
1.00
2.21
1.08
1.15
1.21
1.02
4.52
3.06
7.13
f1 (ppm)

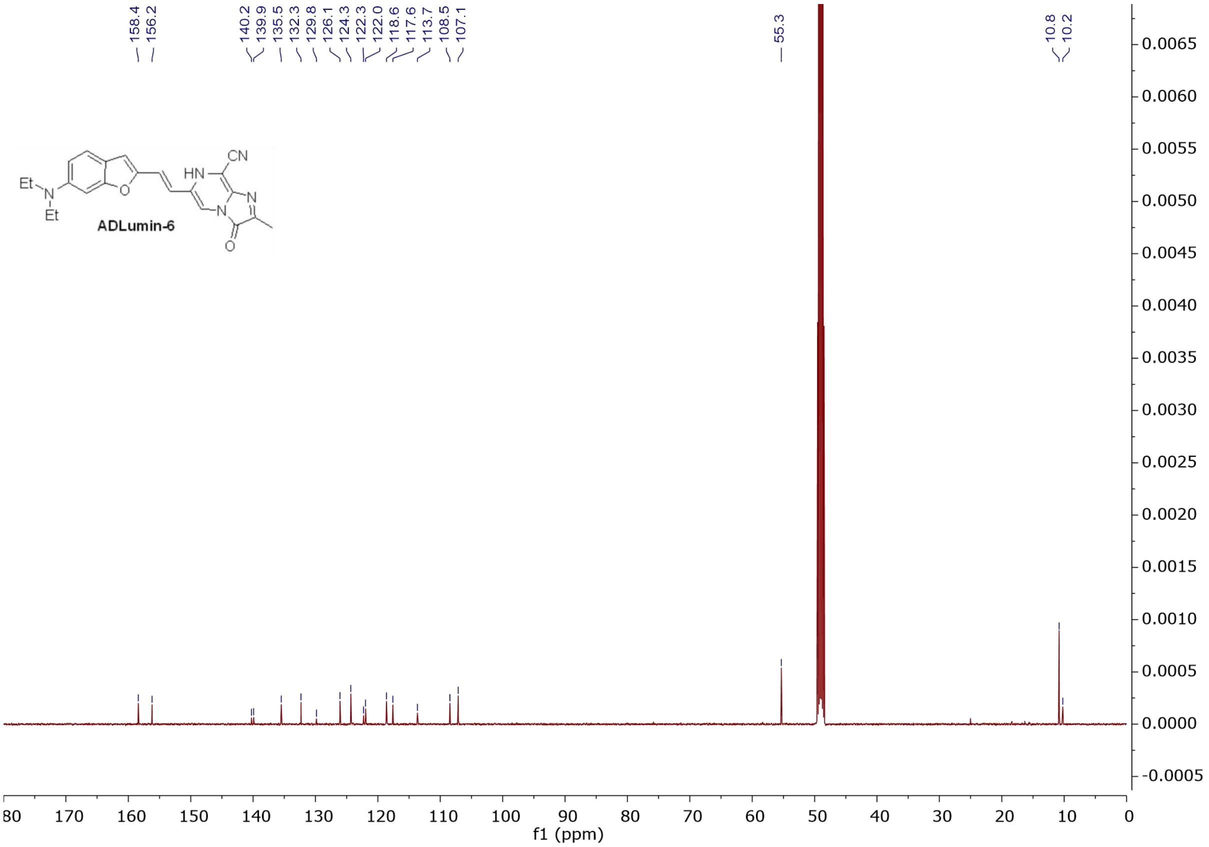
158.4
156.2
140.2
139.9
135.5
132.3
129.8
126.1
124.3
122.3
122.0
118.6
117.6
113.7
108.5
107.1
55.3
10.8
10.2
CN
HN
N
Et
N
Et
O
O
ADLumin-6
f1 (ppm)

Spectrum

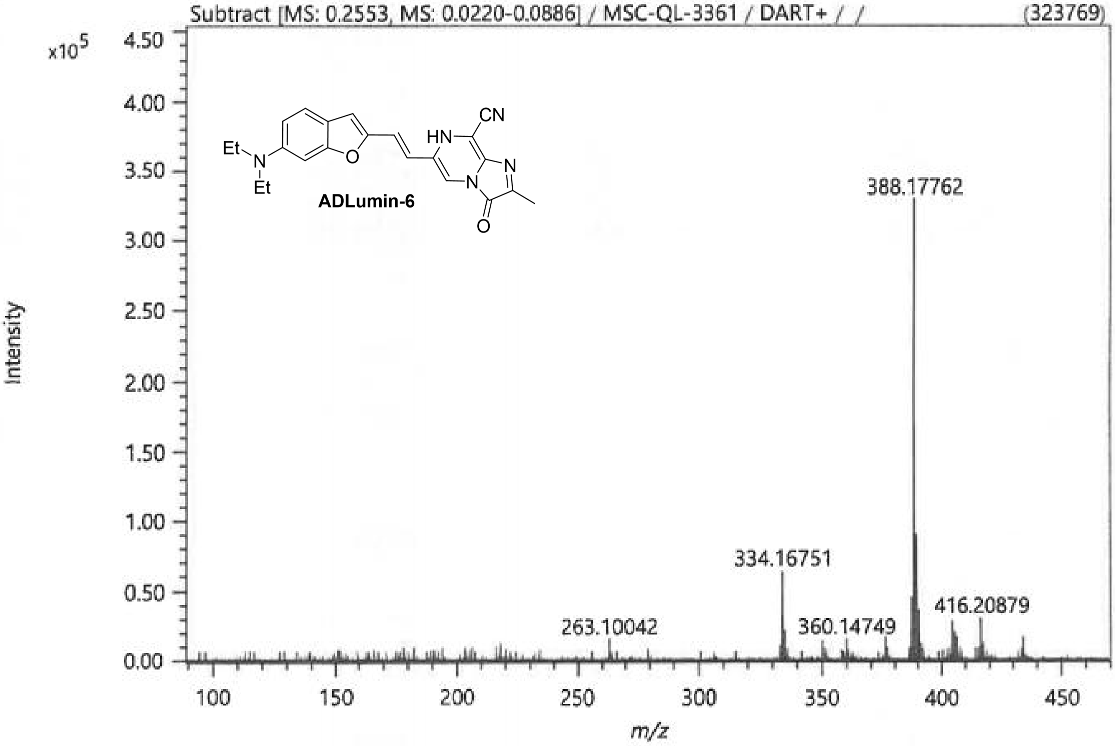


Elemental Composition

Parameters

| | |
|---|---|
| Tolerance: | 6.00 ppm |
| Electron: | Odd/Even |
| Charge: | +1 |
| DBE: | -50.5 - 150.0 |

Elements Set 1:

| Symbol | C | H | O | N | S | Br |
|---|---|---|---|---|---|---|
| Min | 1 | 1 | 1 | 1 | 0 | 0 |
| Max | 100 | 100 | 2 | 5 | 0 | 0 |

Results

| Mass | Intensity | Formula | Calculated Mass | Mass Difference [ppm] | DBE |
|---|---|---|---|---|---|
| 388.17762 | 323768.56 | C22 H22 N5 O2 | 388.17680 | 2.12 | 14.5 |

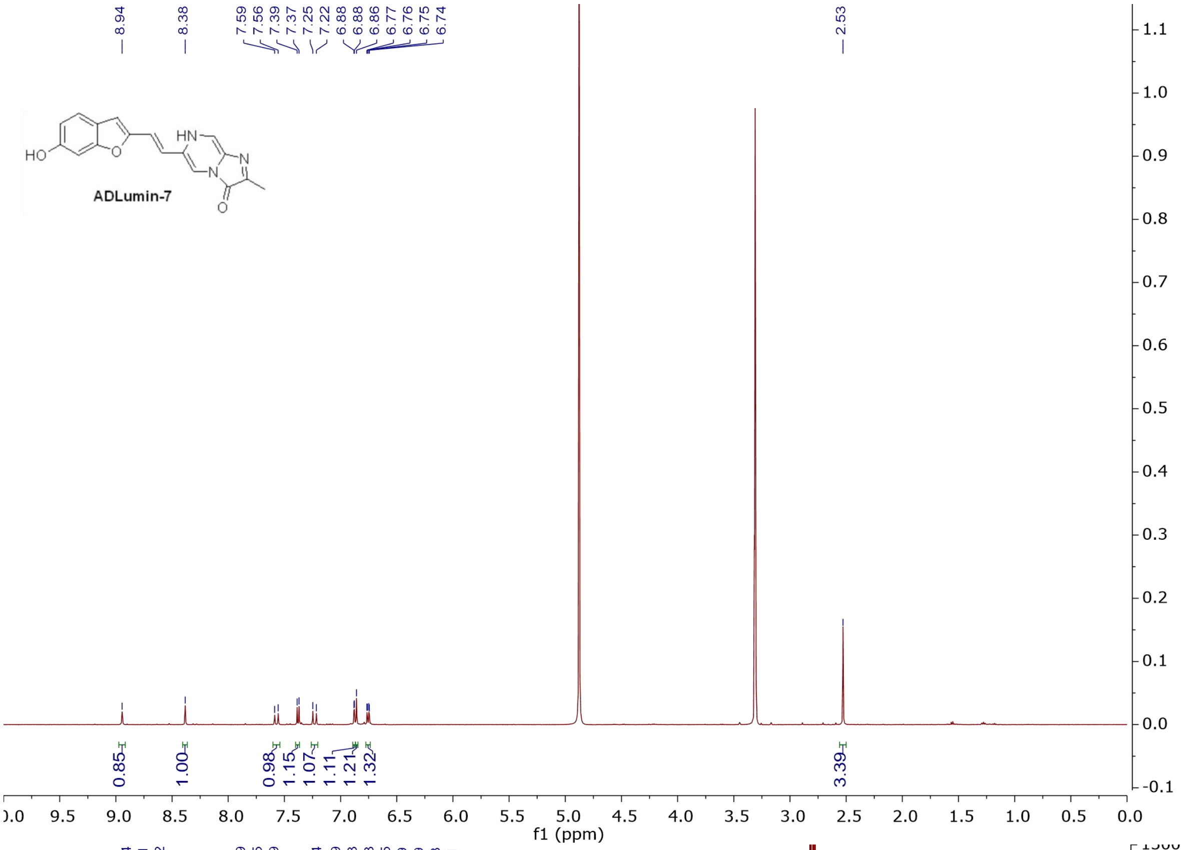
8.94
8.38
7.59
7.56
7.39
7.37
7.25
7.22
6.88
6.88
6.86
6.77
6.76
6.75
6.74
2.53
HO
HN
N
N
O
ADLumin-7
0.85
1.00
0.98
1.15
1.07
1.11
1.21
1.32
3.39
f1 (ppm)

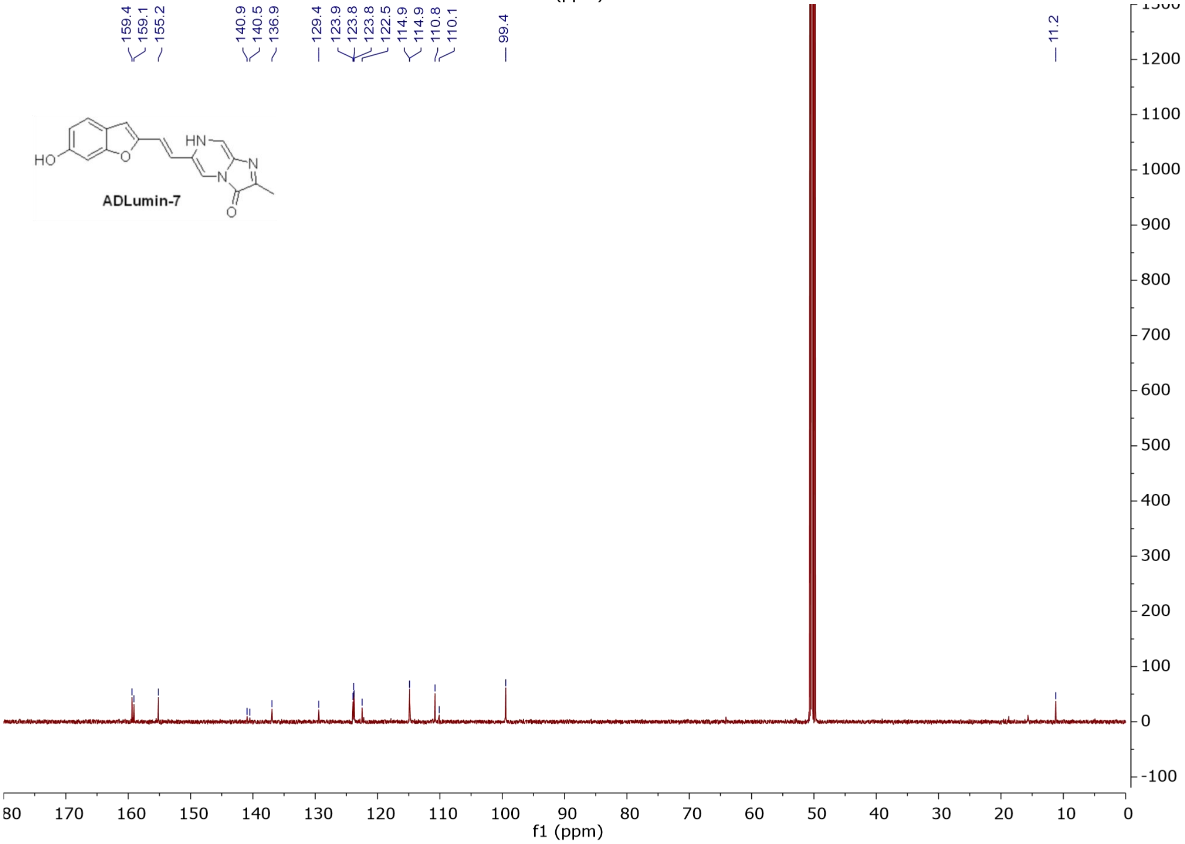
159.4
159.1
155.2
140.9
140.5
136.9
129.4
123.9
123.8
123.8
122.5
114.9
114.9
110.8
110.1
99.4
11.2
HO
HN
N
N
O
ADLumin-7
f1 (ppm)

Spectrum

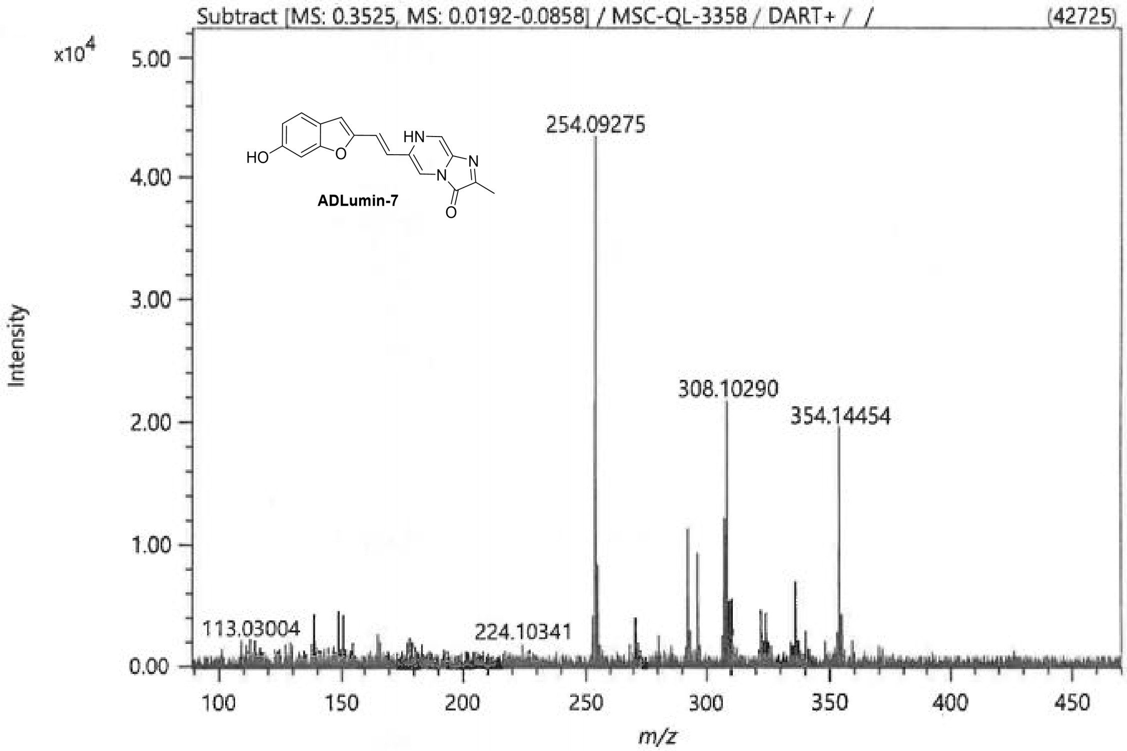


Elemental Composition

Parameters

| | |
|---|---|
| Tolerance: | 6.00 ppm |
| Electron: | Odd/Even |
| Charge: | +1 |
| DBE: | -50.5 - 150.0 |

Elements Set 1:

| Symbol | C | H | O | N | S | Br |
|---|---|---|---|---|---|---|
| Min | 1 | 1 | 1 | 1 | 0 | 0 |
| Max | 100 | 100 | 3 | 3 | 0 | 0 |

Results

| Mass | Intensity | Formula | Calculated Mass | Mass Difference [ppm] | DBE |
|---|---|---|---|---|---|
| 308.10290 | 20981.13 | C17 H14 N3 O3 | 308.10297 | -0.23 | 12.5 |

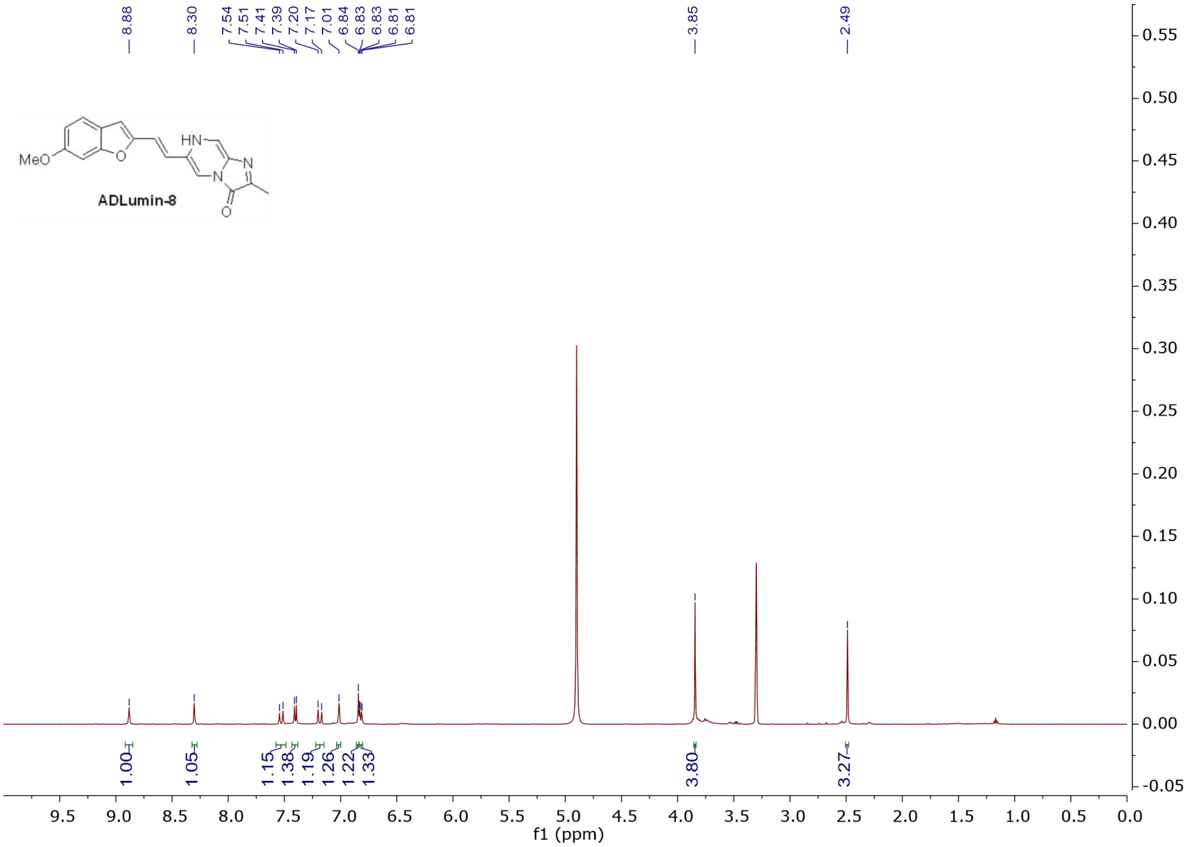

8.88
8.30
7.54
7.51
7.41
7.39
7.20
7.17
7.01
6.84
6.83
6.83
6.81
6.81
3.85
2.49
MeO
O
HN
N
N
O
ADLumin-8
1.00
1.05
1.15
1.38
1.19
1.26
1.22
1.33
3.80
3.27
f1 (ppm)

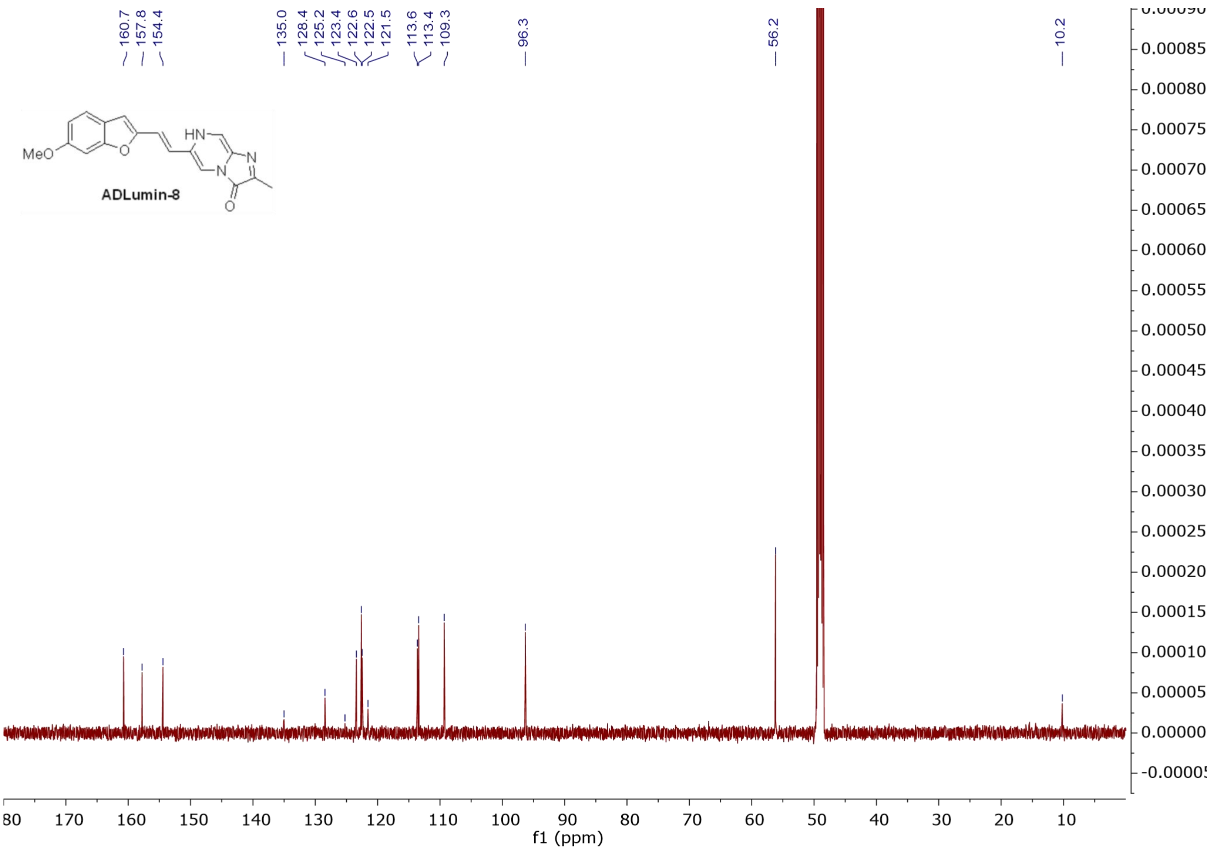
160.7
157.8
154.4
135.0
128.4
125.2
123.4
122.6
122.5
121.5
113.6
113.4
109.3
96.3
56.2
10.2
MeO
O
HN
N
N
O
ADLumin-8
f1 (ppm)

Spectrum

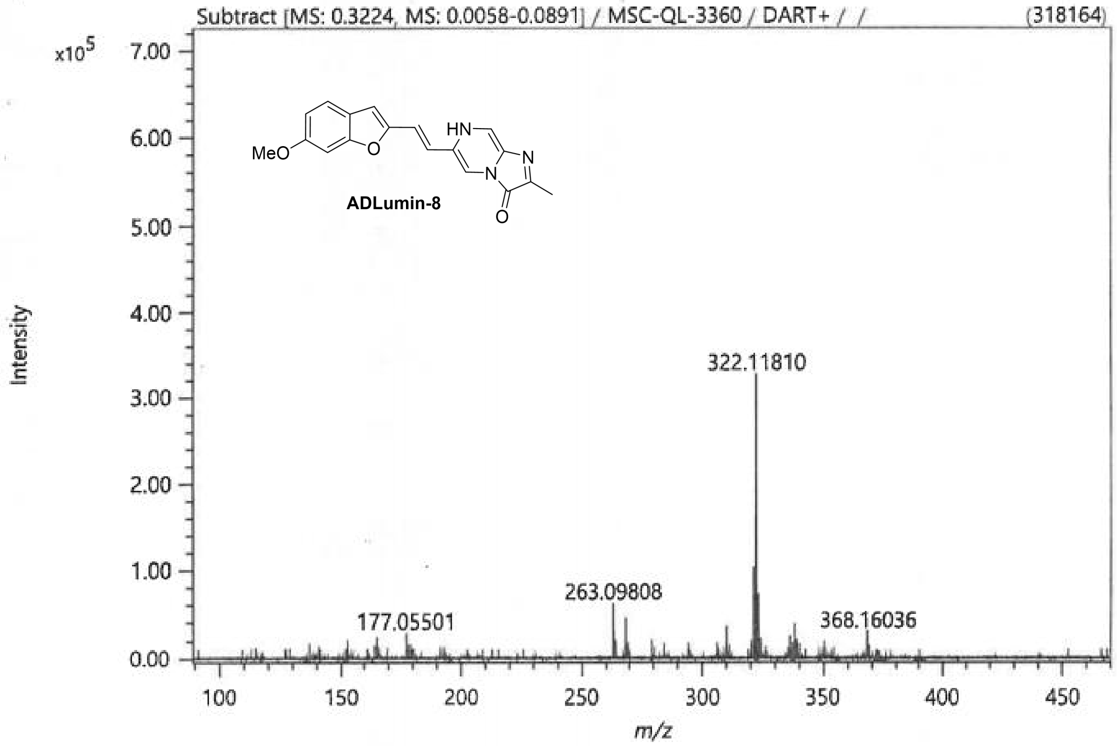


Elemental Composition

Parameters

| | |
|---|---|
| Tolerance: | 6.00 ppm |
| Electron: | Odd/Even |
| Charge: | +1 |
| DBE: | -50.5 - 150.0 |

Elements Set 1:

| Symbol | C | H | O | N | S | Br |
|---|---|---|---|---|---|---|
| Min | 1 | 1 | 1 | 1 | 0 | 0 |
| Max | 100 | 100 | 3 | 3 | 0 | 0 |

Results

| Mass | Intensity | Formula | Calculated Mass | Mass Difference [ppm] | DBE |
|---|---|---|---|---|---|
| 322.11810 | 318164.07 | C18 H16 N3 O3 | 322.11862 | -1.61 | 12.5 |

**SI References**

Sample References:


1. C. Ran, X. Xu, S. B. Raymond, B. J. Ferrara, K. Neal, B. J. Bacskai, Z. Medarova, A. Moore, Design, Synthesis, and Testing of Difluoroboron-Derivatized Curcumins as Near-Infrared Probes for in Vivo Detection of Amyloid-$\beta$ Deposits. *J. Am. Chem. Soc.* 131, 15257-15261 (2009).
2. Y. Y. Zhao, L. Zhang, Z. X. Chen, B. Y. Zheng, M. R. Ke, X. S. Li, J. D. Huang, Nanostructured Phthalocyanine Assemblies with Efficient Synergistic Effect of Type I Photoreaction and Photothermal Action to Overcome Tumor Hypoxia in Photodynamic Therapy. *J. Am. Chem. Soc.* 143, 13980-13989 (2021).
3. E. Epifanovsky, A. T. B. Gilbert, X. Feng, J. Lee, Y. Mao, N. Mardirossian, P. Pokhilko, A. F. White, M. P. Coons, A. L. Dempwolff, Z. Gan, D. Hait, P. R. Horn, L. D. Jacobson, I. Kaliman, J. Kussmann, A. W. Lange, K. U. Lao, D. S. Levine, J. Liu, S. C. McKenzie, A. F. Morrison, K. D. Nanda, F. Plasser, D. R. Rehn, M. L. Vidal, Z.-Q. You, Y. Zhu, B. Alam, B. J. Albrecht, A. Aldossary, E. Alguire, J. H. Andersen, V. Athavale, D. Barton, K. Begam, A. Behn, N. Bellonzi, Y. A. Bernard, E. J. Berquist, H. G. A. Burton, A. Carreras, K. Carter-Fenk, R. Chakraborty, A. D. Chien, K. D. Closser, V. Cofer-Shabica, S. Dasgupta, M. de Wergifosse, J. Deng, M. Diedenhofen, H. Do, S. Ehlert, P.-T. Fang, S. Fatehi, Q. Feng, T. Friedhoff, J. Gayvert, Q. Ge, G. Gidofalvi, M. Goldey, J. Gomes, C. E. González-Espinoza, S. Gulania, A. O. Gunina, M. W. D. Hanson-Heine, P. H. P. Harbach, A. Hauser, M. F. Herbst, M. Hernández Vera, M. Hodecker, Z. C. Holden, S. Houck, X. Huang, K. Hui, B. C. Huynh, M. Ivanov, Á. Jász, H. Ji, H. Jiang, B. Kaduk, S. Kähler, K. Khistyaev, J. Kim, G. Kis, P. Klunzinger, Z. Koczor-Benda, J. H. Koh, D. Kosenkov, L. Koulias, T. Kowalczyk, C. M. Krauter, K. Kue, A. Kunitsa, T. Kus, I. Ladjánszki, A. Landau, K. V. Lawler, D. Lefrancois, Lehtola, Li, R. R.; Li, Y.-P.; Liang, J.; Liebenthal, M.; Lin, H.-H.; Lin, Y.-S.; Liu, F.; Liu, S. K.-Y.; M. Loipersberger, A. Luenser, A. Manjanath, P. Manohar, E. Mansoor, S. F. Manzer, S.-P. Mao, A. V. Marenich, T. Markovich, S. Mason, S. A. Maurer, P. F. McLaughlin, M. F. S. J. Menger, J.-M. Mewes, S. A. Mewes, P. Morgante, J. W. Mullinax, K. J. Oosterbaan, G. Paran, A. C. Paul, S. K. Paul, F. Pavošević, Z. Pei, S. Prager, E. I. Proynov, Á. Rák, E. Ramos-Cordoba, B. Rana, A. E. Rask, A. Rettig, R. M. Richard, F. Rob, E. Rossomme, T. Scheele, M. Scheurer, M. Schneider, N. Sergueev, S. M. Sharada, W. Skomorowski, D. W. Small, C. J. Stein, Y.-C. Su, E. J. Sundstrom, Z. Tao, J. Thirman, G. J. Tornai, T. Tsuchimochi, N. M. Tubman, S. P. Veccham, O. Vydrov, J. Wenzel, J. Witte, A. Yamada, K. Yao, S. Yeganeh, S. R. Yost, A. Zech, I. Y. Zhang, X. Zhang, Y. Zhang, D. Zuev, A. Aspuru-Guzik, A. T. Bell, N. A. Besley, K. B. Bravaya, B. R. Brooks, D. Casanova, J.-D. Chai, S. Coriani, C. J. Cramer, G. Cserey, A. E. DePrince, III; R. A. DiStasio, Jr.; A. Dreuw, B. D. Dunietz, T. R. Furlani, W. A. Goddard, III; S. Hammes-Schiffer, T. Head-Gordon, W. J. Hehre, C.-P. Hsu, T.-C. Jagau, Y. Jung, A. Klamt, J. Kong, D. S. Lambrecht, W. Liang, N. J. Mayhall, C. W. McCurdy, J. B. Neaton, C. Ochsenfeld, J. A. Parkhill, R. Peverati, V. A. Rassolov, Y. Shao, L. V. Slipchenko, T. Stauch, R. P. Steele, J. E. Subotnik, A. J. W. Thom, A. Tkatchenko, D. G. Truhlar, T. Van Voorhis, T. A. Wesolowski, K. B. Whaley, H. L. Woodcock, III; P. M. Zimmerman, S. Faraji, P. M. W. Gill, M. Head-Gordon, J. M. Herbert, A. I. Krylov, Software for the Frontiers of Quantum Chemistry: An Overview of Developments in the Q-Chem 5 Package. *J. Chem. Phys.* 155, 084801 (2021).
4. V. Barone, M. Cossi, Quantum Calculation of Molecular Energies and Energy Gradients in Solution by a Conductor Solvent Model. *J. Phys. Chem. A* 102, 1995-2001 (1998).
5. J.-D. Chai, M. Head-Gordon, Long-Range Corrected Hybrid Density Functionals with Damped Atom–Atom Dispersion Corrections. *Phys. Chem. Chem. Phys.* 10, 6615-6620 (2008).
6. G. W. Spitznagel, T. Clark, J. Chandrasekhar, P. V. R. Schleyer, Stabilization of Methyl Anions by First-Row Substituents. The Superiority of Diffuse Function-Augmented Basis Sets for Anion Calculations. *J. Comput. Chem.* 3, 363-371 (1982).

7. M. C. Davis, T. J. Groshens, D. A. Parrish, Preparation of Cyan Dyes from 6-Diethylaminobenzo[b]furan-2-carboxaldehyde. *Synthetic Commun.* 40, 3008-3020 (2010).
8. S. Martín-Santamaría, J. J. Rodríguez, S. de Pascual-Teresa, S. Gordon, M. Bengtsson, I. Garrido-Laguna, B. Rubio-Viqueira, P. P. López-Casas, M. Hidalgo, B. de Pascual-Teresa, A. Ramos, New scaffolds for the design of selective estrogen receptor modulators. *Org. Biomol. Chem.* 6, 3486-3496 (2008).
